# High diversity in the regulatory region of Stx-converting bacteriophage genomes

**DOI:** 10.1101/2021.11.24.469858

**Authors:** Annette Fagerlund, Marina Aspholm, Grzegorz Węgrzyn, Toril Lindbäck

**Affiliations:** Nofima, Norwegian Institute of Food, Fisheries and Aquaculture Research, Ås, Norway; Department of Paraclinical Sciences, Faculty of Veterinary Medicine, Norwegian University of Life Sciences, Ås, Norway; Department of Molecular Biology, Faculty of Biology, University of Gdañsk, Gdañsk, Poland

**Keywords:** EHEC, Stx phage, bacteriophage genetics, lysogen, lytic, phage replication

## Abstract

Shiga toxin (Stx) is the major virulence factor of enterohemorrhagic *Escherichia coli* (EHEC), and the *stx* genes are carried by temperate bacteriophages (Stx phages). The switch between lysogenic and lytic life cycle of the phage, which is crucial for Stx production and for severity of the disease, is regulated by the CI repressor. CI maintain latency by preventing transcription of the replication proteins. Three <u>E</u>HEC phage <u>r</u>eplication <u>u</u>nits (Eru1-3) in addition to the classical lambdoid replication region have been described previously, and Stx phages carrying the Eru1 replication region were associated with highly virulent EHEC strains. In this study, we have classified the Eru replication region of 419 Stx phages. In addition to the lambdoid replication region and the three already described Erus, ten novel Erus (named Eru4 to Eru13) were detected. The lambdoid type, Eru1, Eru4 and Eru7 seem to be widely distributed in Western Europe. Notably, EHEC strains involved in severe outbreaks in England and Norway carry Stx phages with Eru1, Eru2, Eru5 and Eru7 replication regions. Phylogenetic analysis of CI repressors from Stx phages revealed eight major clades that largely separate according to Eru type. The classification of replication regions and CI proteins of Stx phages provides an important platform for further studies aimed to assess how characteristics of the replication region influence the regulation of phage life cycle and, consequently, the virulence potential of the host EHEC strain.

**IMPORTANCE:** EHEC is an emerging health challenge worldwide and outbreaks caused by this pathogen tend to be more frequent and severe. Increased knowledge on how characteristics of the replication region influence the virulence of *E. coli* may be used for more precise identification of high-risk EHEC strains.

## INTRODUCTION

Enterohemorrhagic *Escherichia coli* (EHEC) is an important foodborne pathogen, responsible for disease in humans ranging from uncomplicated diarrhea to severe conditions such as hemorrhagic colitis and hemolytic uremic syndrome (HUS). EHEC is an emerging health challenge worldwide and outbreaks caused by this pathogen tend to become more frequent and severe: WHO has estimated that 10% of patients with EHEC infection develop HUS. Since the first major EHEC outbreak in 1982, caused by serotype O157:H7, the world has experienced multiple outbreaks of EHEC disease involving other serotypes than O157:H7 and new variants are constantly emerging (1, 2). Shiga toxin (Stx) is the major virulence factor of EHEC, and it exists in two distinct forms, Stx1 and Stx2. Each form comprises several subtypes (3) where some subtypes such as Stx2a are associated with severe disease while Stx2c is considered less potent (4, 5).

The genes encoding Stx are carried by temperate bacteriophages (Stx phages). After infection, Stx phages follow either a lysogenic or lytic pathway. The lysogenic pathway involves integration of phage DNA into the host genome and replication of the phage genetic material along with the chromosome of the host cell. The lytic pathway leads to proliferation of the Stx phage, death of the host bacterial cell and release of new phage particles (6). The proliferation of Stx phages is also accompanied by production and release of substantial amounts of Stx toxin (7). As the amount of produced Stx influences the severity of the disease, the mechanisms regulating the switch from lysogenic to lytic life cycle is highly relevant for the pathogenicity of the host *E. coli* strain.

Since the first sequenced Stx phages shared substantial genomic similarity to phage lambda it has been assumed that they behave similarly (8, 9). The increasing availability of whole genome sequences has revealed that Stx-encoding prophages are very diverse and, sometimes, exhibit only limited similarity towards phage lambda (10). We have previously reported Stx phages with non-lambdoid replication regions and named the regions Eru (<u>E</u>HEC phage <u>r</u>eplication <u>u</u>nit) (10). The non-lambdoid Stx phages completely lack the *O* and *P* genes, encoding proteins involved in replication initiation of the lambdoid phage genome, and instead carry genes which have previously not been described in connection to replication of Stx phages. Three non-lambdoid Stx phage replications, Eru1-3, have so far been described (10). One of the Eru types, Eru1, is carried by the highly pathogenic EHEC strains that caused the Norwegian O103:H25 outbreak in 2006 and the large O104:H4 outbreak in Europe in 2011. It was also shown that Eru1 phages exhibited a less stable lysogenic state than the classical lambdoid Stx phages, which could increase the pathogenicity of the host *E. coli* strain (10). The majority of EHEC strains carrying Eru1, Eru2 and Eru3 type of Stx phages were US isolates whose genome sequences were submitted to NCBI databases by the United State Department of Agriculture, the US Food and Drug Administration, and the Food-borne Pathogen Omics Research Center.

Despite the high genetic diversity among Stx phage genomes, the phage replication region and the lysis-lysogeny regulatory systems are always located upstream and in the vicinity to the *stx* gene (11). This region mediates the switch between repression and induction of the prophage, and the mechanisms regulating these events have been studied in detail in phage lambda. The key elements responsible for regulating the life cycle of phage lambda are the gene encoding repressor CI (*cI*), the promoter binding the CI repressor and the adjacent upstream genes, transcribed in the opposite direction of *cI* (Fig. 1) (12, 13). The lambda CI repressor downregulates expression of genes involved in production of new phage particles, i.e., the lytic cycle, by specific binding to the promoter region of the adjacent genes encoding the O and P proteins which initiate replication of the lambda genome (14). The crystal structure of CI has been solved and revealed that the protein is functional as a homodimer and that repression occurs when two subunits bind cooperatively to adjacent operator sites on the DNA (15). The C-terminal domain mediates the dimer formation and the dimer-dimer interactions enable CI to bind cooperatively to two or more operator sites (16, 17) while the N-terminal domain contains a helix-turn-helix DNA-binding domain (18, 19). Upon DNA damage, the SOS-response protein RecA becomes activated and may in lysogenic cells stimulate autocleavage of CI (20). Cleaved CI can no longer bind to DNA and repression of the promoters in the replication module is thus relieved.

**Figure 1:**
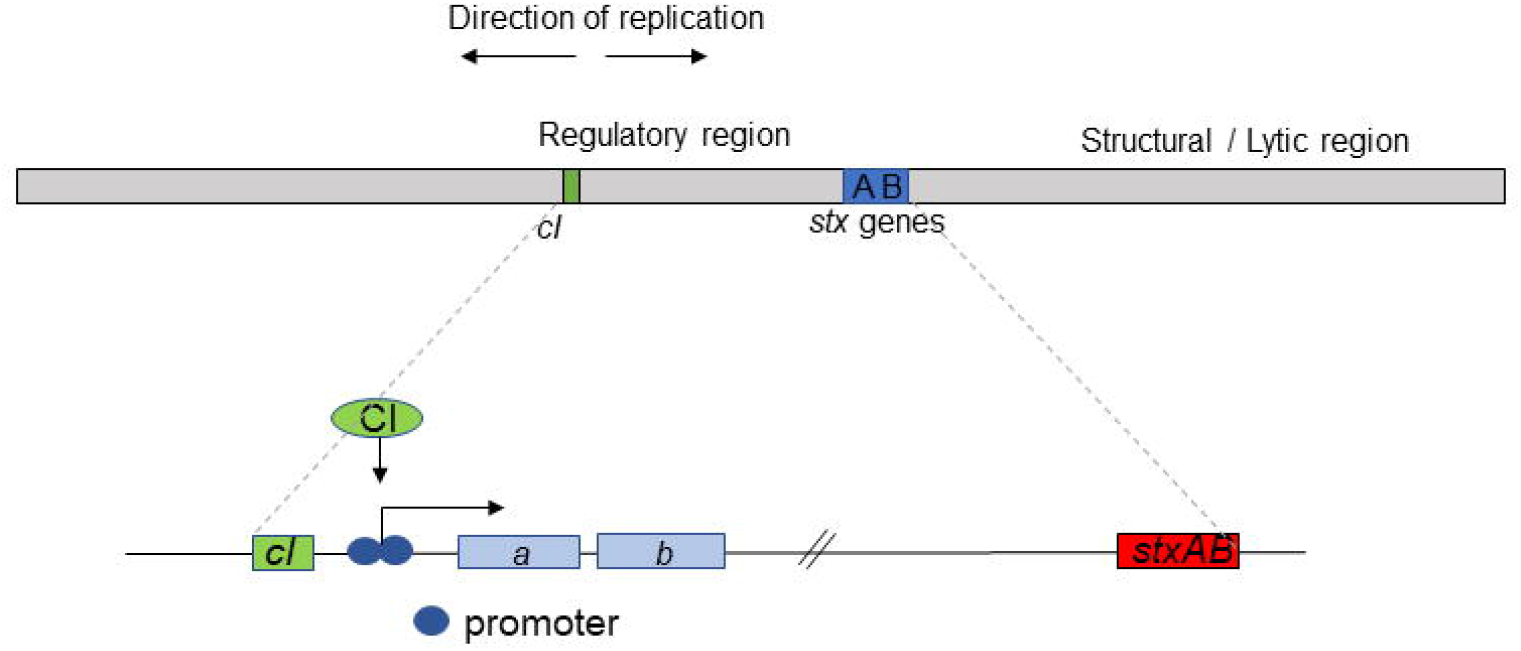
A schematic overview of the genome of an Stx phage. The boxes labeled *a* and *b* indicate the replication genes which are represented by *O* and *P* in phage lambda and by other less characterized genes in Erul-3 (10).

Some EHEC strains appear more virulent than others and the type of Stx produced is known to contribute significantly to the pathogenicity of the EHEC strain (5) but the amount of toxin produced should also be considered. The increasing number of outbreaks of gastrointestinal disease and HUS caused by EHEC have stimulated studies on the Stx phages to better understand their contribution to the pathogenicity of the host *E. coli* strain. However, there is still very limited knowledge on how the different types of replication regions seen among the Stx phages influences the stability of the lysogenic state and the switch to lytic cycle. In this study, we have classified the CI repressor sequences of 260 Stx phages into clade I-VIII and their replication regions into 13 Eru types to provide a platform for further studies of how the genetic structure of the Stx prophages influences the virulence potential of the host EHEC strain.

## RESULTS

Eru types were defined by the identity of the proteins encoded by the two genes located directly upstream and in opposite direction of *cI*. Four novel Eru phage types (Eru4-7) were detected among 120 Stx-converting phage genomes retrieved from NCBI virus database (Fig. 2;Table S1 in supplemental material), while an additional six novel Eru types, Eru8 to Eru13, were detected among 299 genome sequences obtained from ten examined NCBI BioProjects (Fig. 2; Table S2 in supplemental material). Eru2 and Eru3, described in a previous study, both carry genes encoding a protein of unknown function and a helicase directly upstream of *cI* (10). However, since the two unknown proteins share a low sequence identity (10%) phages carrying these protein combinations were still assigned to different Eru types (10). Phages representing each Eru type is listed in Table 1 as reference phages for each Eru type. The distribution of Eru types found among the 120 sequenced Stx phages are shown in Table 2. The relatively high number of phage genomes belonging to Eru2, Eru3 and Eru7 could be due to a bias related to the number of deposited sequences from different studies (Table S1 in supplemental material).

**Figure 2:**
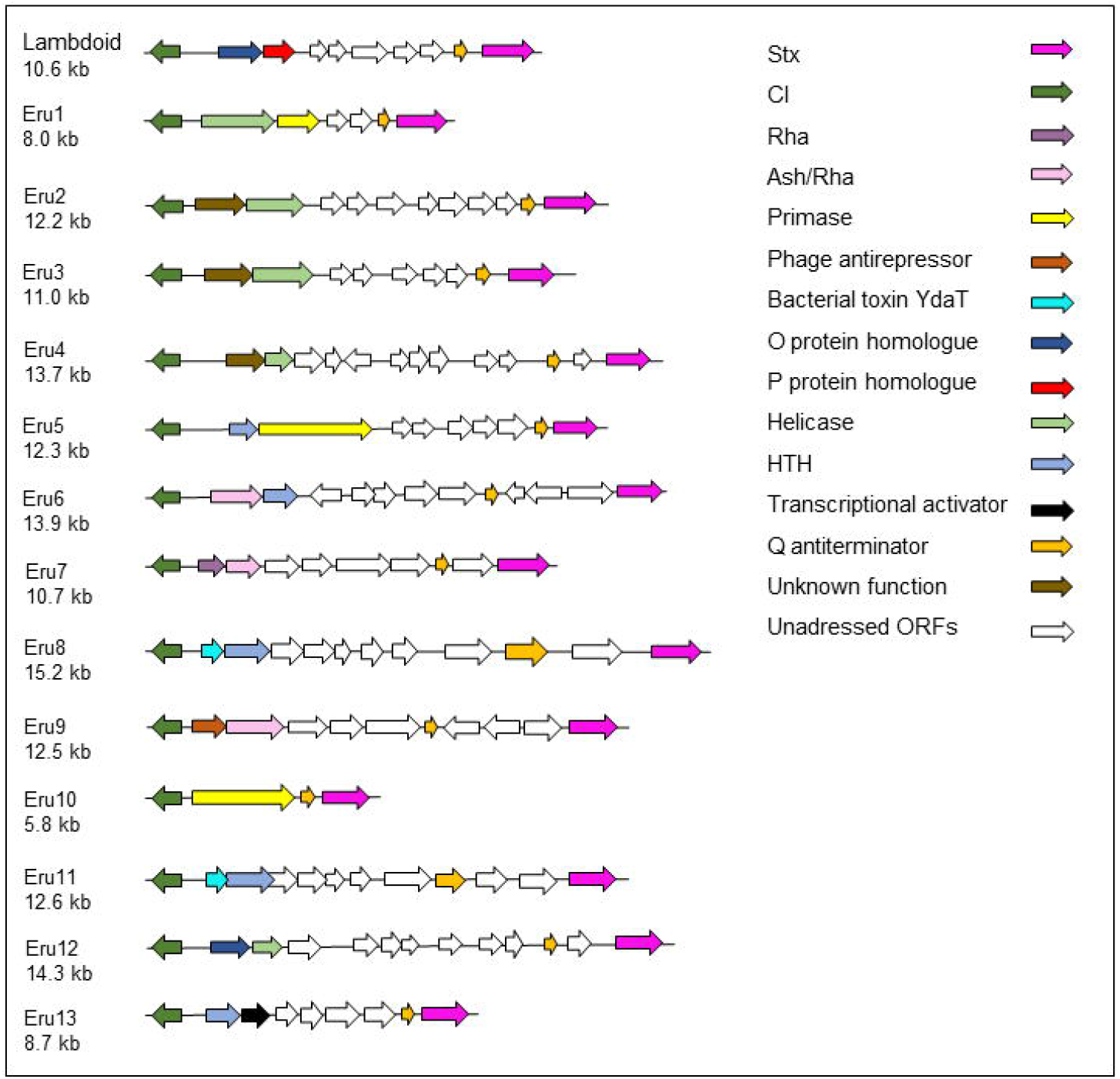
Physical maps of the region between *cI* (green) and *stx* (pink). The color code also indicates the putative function of the proteins encoded by the genes directly upstream of *cI*. White arrows indicate open reading frames (ORFs) which are not addressed in this study.

**Table 1:**
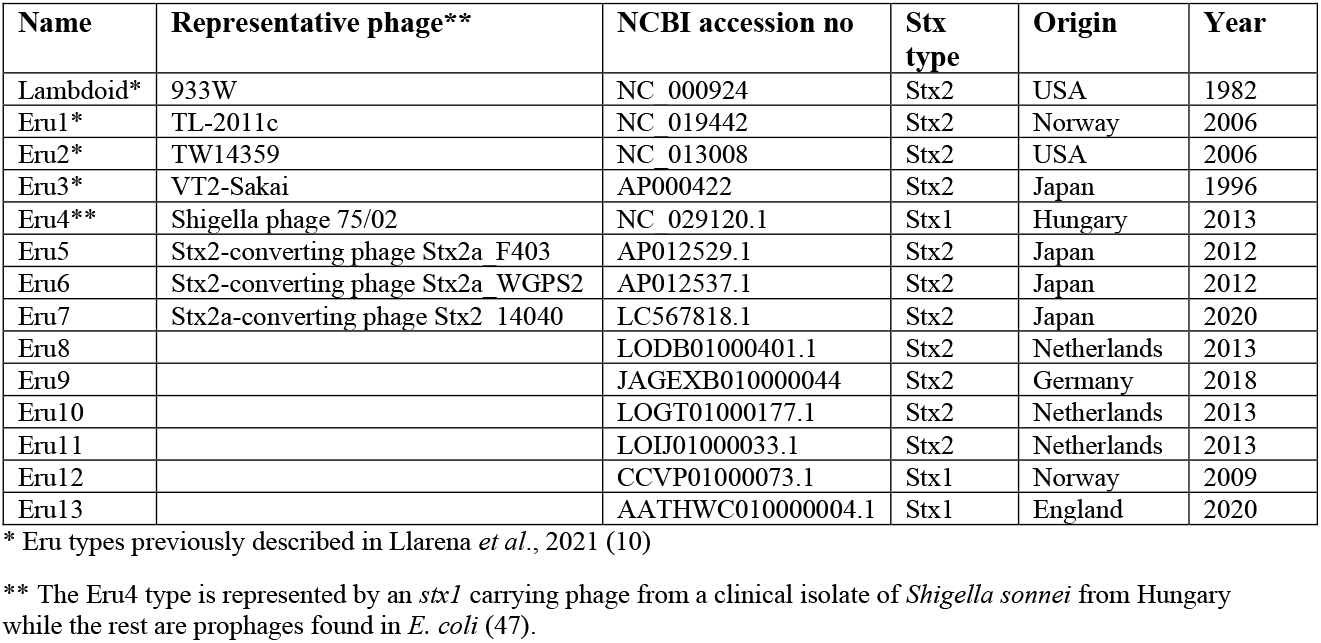
Accession numbers of sequences representing each Eru type

| Name | Representative phage** | NCBI accession no | Stx type | Origin | Year |
| --- | --- | --- | --- | --- | --- |
| Lambdoid* | 933W | NC_000924 | Stx2 | USA | 1982 |
| Eru1* | TL-2011c | NC_019442 | Stx2 | Norway | 2006 |
| Eru2* | TW14359 | NC_013008 | Stx2 | USA | 2006 |
| Eru3* | VT2-Sakai | AP000422 | Stx2 | Japan | 1996 |
| Eru4** | Shigella phage 75/02 | NC_029120.1 | Stx1 | Hungary | 2013 |
| Eru5 | Stx2-converting phage Stx2a_F403 | AP012529.1 | Stx2 | Japan | 2012 |
| Eru6 | Stx2-converting phage Stx2a_WGPS2 | AP012537.1 | Stx2 | Japan | 2012 |
| Eru7 | Stx2a-converting phage Stx2_14040 | LC567818.1 | Stx2 | Japan | 2020 |
| Eru8 |  | LODB01000401.1 | Stx2 | Netherlands | 2013 |
| Eru9 |  | JAGEXB010000044 | Stx2 | Germany | 2018 |
| Eru10 |  | LOGT01000177.1 | Stx2 | Netherlands | 2013 |
| Eru11 |  | LOIJ01000033.1 | Stx2 | Netherlands | 2013 |
| Eru12 |  | CCVP01000073.1 | Stx1 | Norway | 2009 |
| Eru13 |  | AATHWC010000004.1 | Stx1 | England | 2020 |

\* Eru types previously described in Llarena *et al.*, 2021 (10)
\*\* The Eru4 type is represented by an *stx1* carrying phage from a clinical isolate of *Shigella sonnei* from Hungary while the rest are prophages found in *E. coli* (47).

**Table 2:** Number of Eru types in the data set of 120 Stx-converting phage genomes retrieved from the NCBI virus database (taxid:10239).

| Eru type | Number | Reference |
| --- | --- | --- |
| lambdoid | 9 | (10) |
| Eru1 | 7 | (10) |
| Eru2 | 57 | (10) |
| Eru3 | 18 | (10) |
| Eru4 | 2 | This study |
| Eru5 | 4 | This study |
| Eru6 | 3 | This study |
| Eru7 | 20 | This study |

### Distribution of Eru types among Stx phages from Western Europe

The national distribution of Eru types found among 299 identified contigs carrying both *stx* and *cI* from ten European BioProjects is shown in Table 3. The distribution of Eru types indicates that the lambdoid and the Eru1, 4 and 7 phage types are among the most common types of Stx phages in Europe, and that Eru7 appears to be particularly widespread.

**Table 3:**
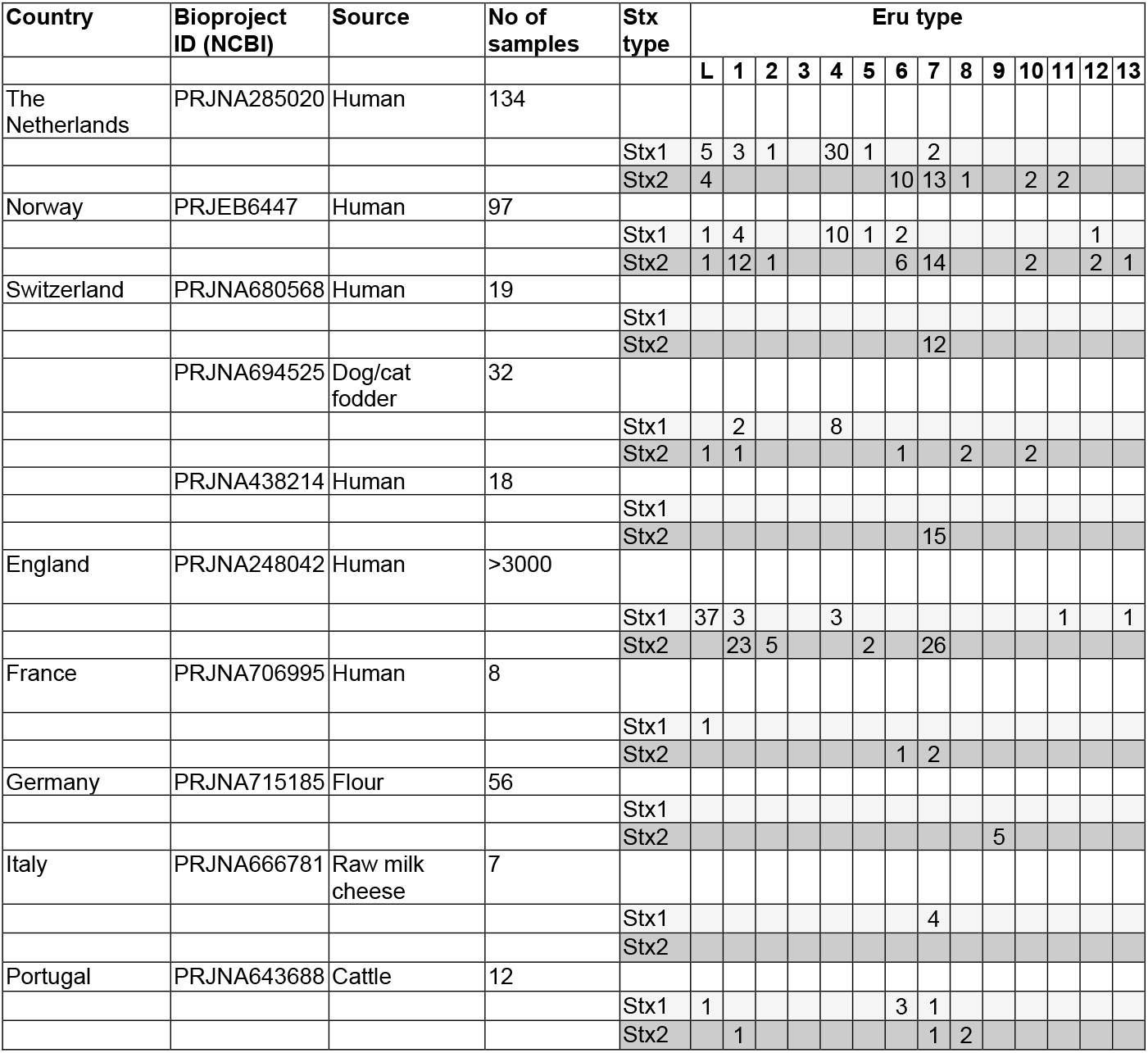
Distribution of the thirteen Eru types (1-13) and the lambdoid (L) type in ten European BioProjects

### The Eru proteins

All Eru phages carry genes encoding different types of DNA binding proteins, such as helicases, primases, or other helix-turn-helix (HTH) motif proteins, in the first and/or second position directly upstream of *cI* (Fig. 2). Eru6, Eru7 and Eru9 phages carry genes encoding proteins of the Phage_pRha protein family (pfam09669) directly upstream of *cI* (Fig. 2). The Rha domain, which contain a winged helix-turn-helix DNA-binding motif, is also found in other temperate phages where it has been suggested to have phage regulatory function (21, 22). Some of the Rha proteins also contain the Ash domain (PF10554), which is present in the ASH protein of bacteriophage P4. However, no function has so far been assigned to this domain (22). Eru4, and the previously described Eru2 and Eru3 (10), encode proteins of unknown function directly upstream of *cI* (Fig. 2). However, there are no similarities between these proteins, and they do not share any previously described protein domains. The primases encoded by genes carried by Eru1, Eru5 and Eru10 phages do not share any sequence similarities (<10 % amino acid identity). The amino acid sequence of the putative helicases encoded by Eru4 and Eru12 are 97% identical and they both share the AAA motif (PF13604) with the Eru1 helicase (10). However, the overall sequence homology between the Eru4 and Eru12 helicases and the Eru1 helicase are low (<10 % amino acid identity).

Genes encoding HTH domain proteins are found in either the first or second position directly upstream of *cI* in Eru5, Eru6, Eru8, Eru11 and Eru13 (Fig. 2). The HTH proteins of Eru5 and Eru6 are 50% identical with a coverage of 66%, the HTH proteins of Eru8 and Eru13 are 59% identical over the total protein sequence, and all five proteins exhibit the HTH_36 motif (PF13730). The HTH proteins of Eru6 and Eru13 also share a motif (PF13814) which is found in protein families essential for relaxation and replication of plasmid DNA (23, 24). Both Eru8 and Eru11 phages carry a gene encoding a protein with homology to the bacterial toxin YdaT (PF06254) directly upstream of *cI*. However, the two Eru-encoded toxin-like proteins share only 34% identity with each other. The shortest distance between *cI* and *stx* was displayed by Eru10, which only carried a bifunctional DNA primase-polymerase motif protein (PF09250) (25) and the Q antiterminator protein (26, 27) in this region. All other Eru phages also carried the gene encoding the antiterminator Q protein between *cI* and *stx*, indicating that this protein is essential for Stx phages.

### Eru types in particularly virulent EHEC

Five different Eru phage types were found among Stx phages from six highly pathogenic EHEC O157:H7 strains that have caused larger outbreaks in the UK (Table 4)(28). Among this panel of phages, all carrying *stx2c* and one carrying *stx2a* are of Eru2 type. Two *stx2a* carrying phages are of the Eru5 type, while the two remaining *stx2a* phages are of types Eru1 and Eru7. The only *stx1a* carrying phage among these isolates has a lambdoid replication region. Among the 97 Norwegian STEC strains in BioProject PRJEB6447, 15 strains caused HUS (29) and 13 of these strains carried *stx2* phages of Eru types 1, 2 or 7.

**Table 4:** Eru type of Stx phages of highly pathogenic STEC O157:H7 isolates from UK

| Strain ID | Reference | NCBI<br>accession no. | Year | Eru type |
| --- | --- | --- | --- | --- |
| E30228 | (48) | VXJO00000000 | 1983 | Stx2a Eru5<br>Stx1a lambdoid |
| E34500 | (49) | VXJN00000000 | 1983 | Stx2a Eru1<br>Stx2c Eru2 |
| E45000 | (28) | VXJM00000000 | 1987 | Stx2a Eru2 |
| E116508 | (28) | VXJP00000000 | 1996 | Stx2a Eru5<br>Stx2c Eru2 |
| 315176 | (50) | VXJQ00000000 | 2014 | Stx2a Eru2 |
| 267849 | (51) | VXJR00000000 | 2016 | Stx2a Eru7<br>Stx2c Eru2 |

### The CI repressors

In order to examine the correspondence between the Eru phage type and the sequence of the CI repressor proteins, a total of 260 annotated CI sequences (Table S3 in supplemental material) were extracted from the phage genomes and used to build a phylogeny (Fig. 3). This analysis grouped the CI proteins into several distinct clades, for which major clades were named I to VIII.

**Figure 3:**
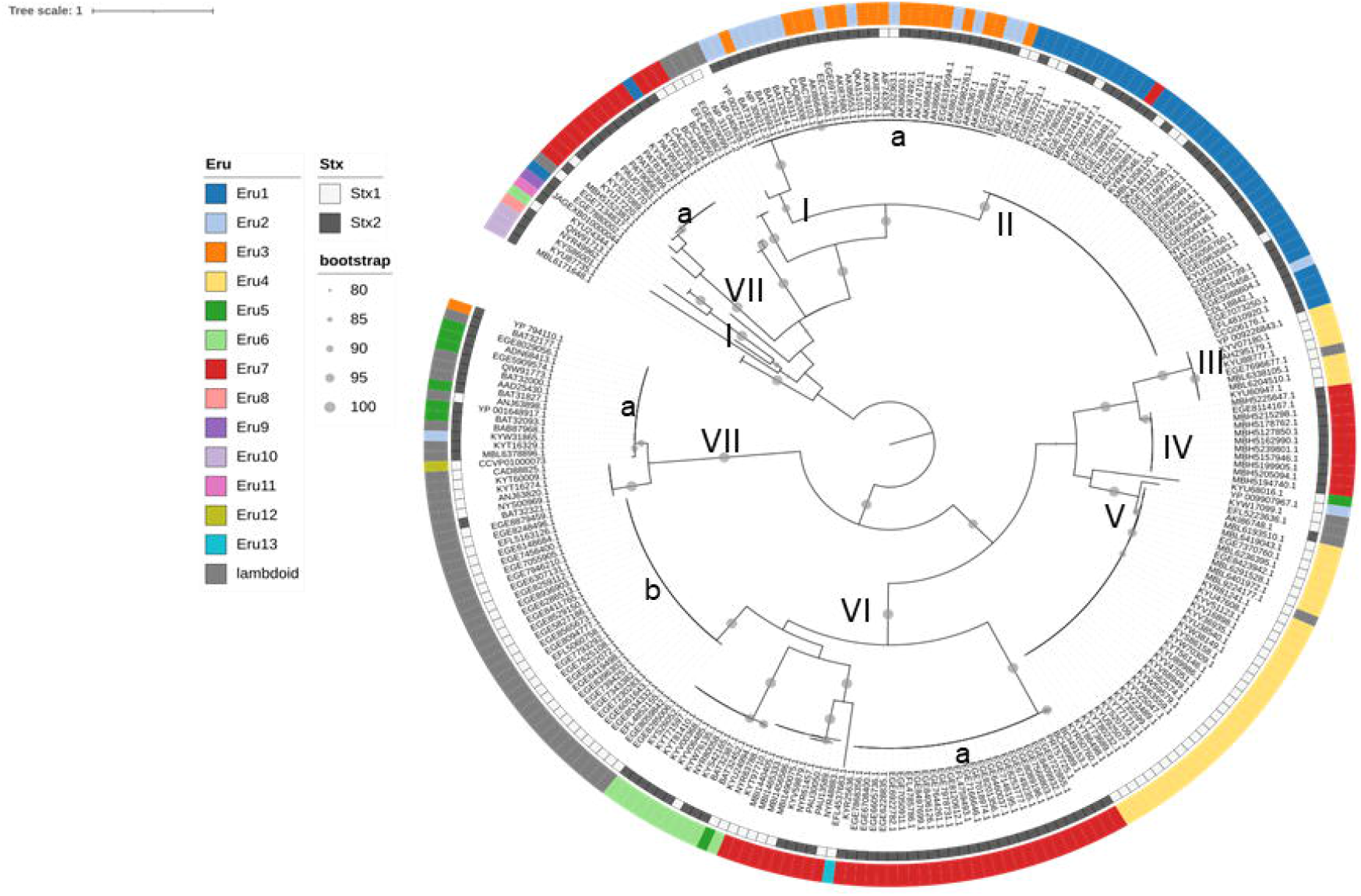
Maximum-likelihood phylogeny of 260 CI protein sequences. The tree was midpoint rooted and bootstrap values >80% are indicated by grey circles. The Stx type is shown in the inner ring and the Eru type is shown in the outer ring. Clades that are discussed in the text are labelled with roman numerals

The CI protein from lambda phage (NP_040628.1) was most closely related to the CI proteins from phages of Eru types 2 and 3, all belonging to Clade I. The CI proteins of Eru2 and Eru3 phages in this clade were all identical and show an overall identity of 61% towards lambda CI. Lambda CI contains two protein domains, a HTH_3 domain (30) and a peptidase_S24 domain, which executes the CI autolysis (31, 32). The two domains are conserved within the CI proteins belonging to Clades I, II, IV, V, VI and VII (Fig. 4). However, the CI proteins of Clade III and YP_009907967.1 in Clade V lack the HTH domain, while Clade VIIIa CI proteins lack the peptidase domain and instead exhibit an additional HTH domain (Fig. 4).

**Figure 4:**
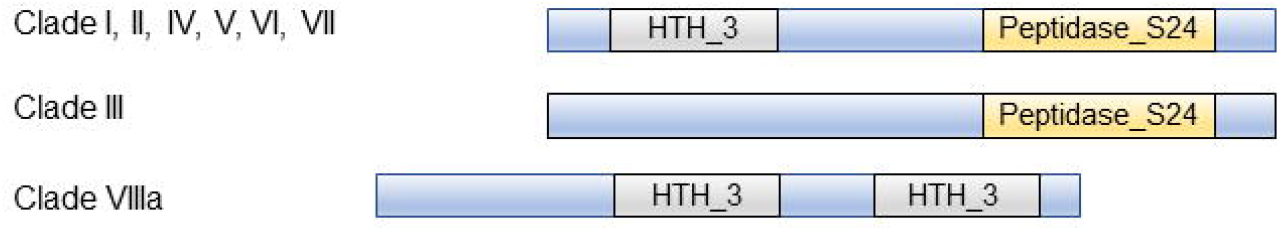
Domain structures of Stx phage CI repressors of Clade I-VIII. HTH_3 domains (grey) and Peptidase_S24 domains (yellow) were assigned according to Pfam.

In contrast to the observed high homology between CI proteins within a clade, the homology between the clades was low (Table S4 in supplemental material). The highest CI homology was seen between Clades I and II (51%) and between Clades III and IV (60%). An amino acid sequence alignment of CI sequences from Clades I to VII is shown in Fig. 5. The alignment revealed six amino acids conserved throughout all clades, one of which was the lambda CI autocleavage residue S150 (16).

**Figure 5:**
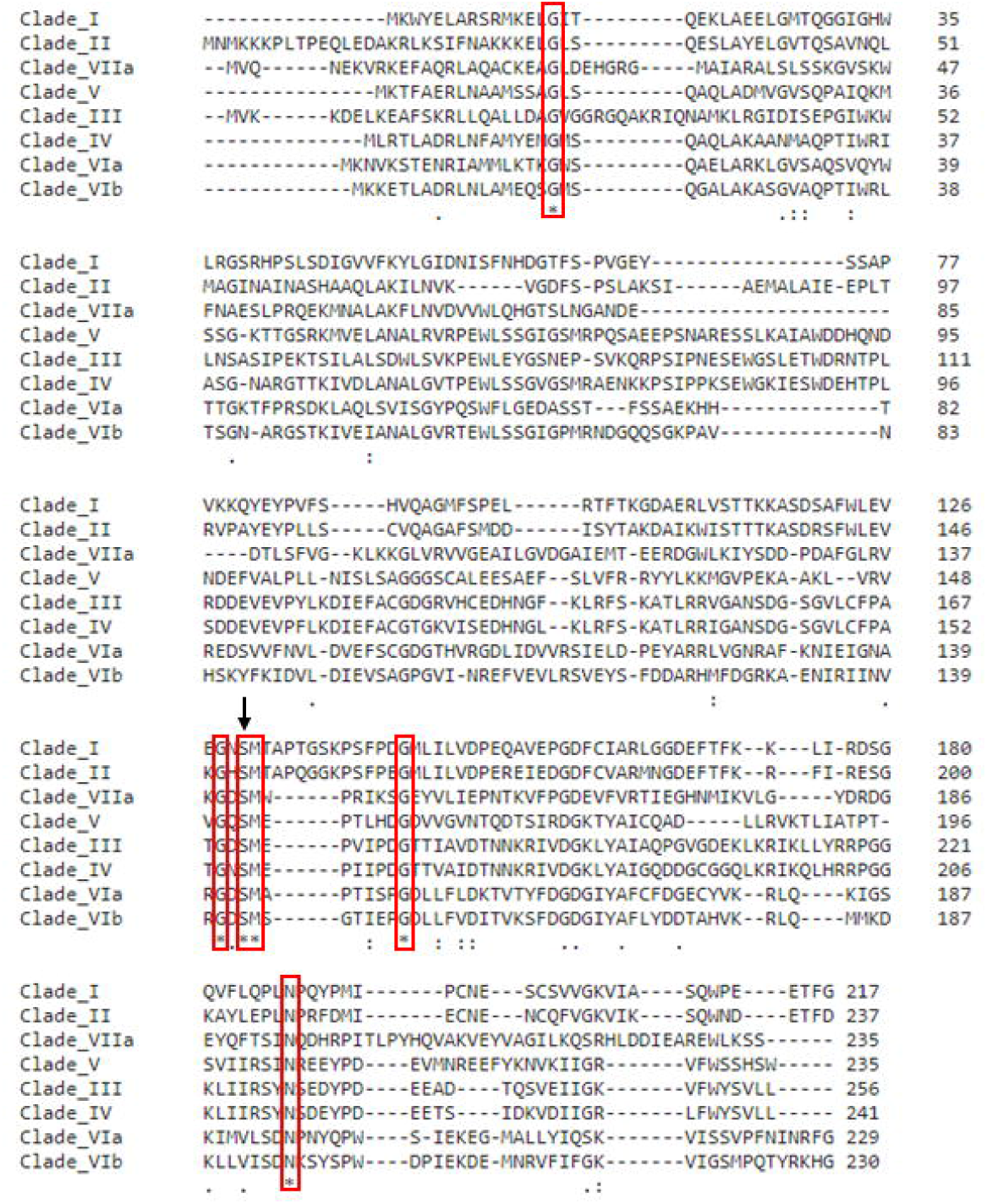
Sequence alignment of Clade I-VII Stx phage CI sequences. CI protein from Clade VIII is not included in the alignment due to large structural differences (see Fig. 4). Red boxes indicate the six amino acids that were conserved throughout all clades and the black arrow indicates the CI autocleavage residue found in this type of repressors (16).

### Strong correlation between CI Clades and Eru type

There is a strong coherence between CI clades and Eru types which is not unexpected in light of their neighboring location in the phage genome. CI proteins belonging to Clades III and V are almost exclusively co-present with Eru4 replication proteins and the lambdoid replication type is mostly found in connection with Clades VIb and VII (Fig. 3). Similarly, the genes encoding CI proteins belonging to Clade II are almost exclusively located directly upstream of the genes defining Eru1, while those belonging to Clade Ia are located upstream of Eru2 and Eru3 (Fig. 3). However, a specific CI clade are not necessarily restricted to a specific Eru type and may regulate expression of different Eru types (Fig. 3). CI proteins of Clades III, V and VIb are linked to the lambdoid or Eru4 types and are mainly found in Stx1 producing phages (Fig. 3).

## DISCUSSION

The present study show that Stx phages are genetically much more diverse than previously anticipated, and this finding is important as differences in phages replication modules has been suggested to influence the stability of the lysogenic state and the pathogenic potential of the host *E. coli* strain (10). The Eru type was here defined based on the type of proteins encoded by the two genes located directly upstream of *cI*. This definition is less differentiating than the definition used by Llarena *et al*. (10) where the entire region between *cI* and *stx* was considered. However, due to the large variation of genes located between *cI* and *stx*, revealed in this study, we found that defining Eru type based on the identity of the two proximal genes set the discrimination level to an appropriate level of sensitivity.

Stx phages have traditionally been classified into the group of lambdoid phages based on similarity in behavior, genetic structure, and regulatory system. In phage lambda and lambdoid Stx phages, the assembly of the replication complex has been studied to detail (33) but there is so far no knowledge about the proteins involved in the replication process of Eru phages. Eru7 seems to be the most widespread Eru type in Europe and Eru7 phages, together with Eru6 and Eru9, encode proteins containing Rha or Rha/Ash domains. Rha domain proteins are common among temperate bacteriophages, prophages, and large eukaryotic DNA viruses and is suggested to function as a regulatory protein that is involved in controlling the switch between lytic and lysogenic lifestyle (34) but very little is known about the function of Ash domain proteins (21, 22, 35). However, none of these proteins have previously been associated with replication of Stx phages and it is of great interest to examine this aspect especially since Eru7 Stx phages seems to be among the more common Eru types.

In phage lambda and lambdoid Stx phages, the CI repressor regulates expression of the *O* and *P* replication genes (13). The *cI* gene is also present in the genomes of Eru phages suggesting that a similar regulatory mechanism is at play in non-lambdoid Stx phages. The genes located directly upstream of *cI* varies extensively between different Eru types, although most of them encode DNA binding proteins such as helicases, primases or other HTH motif proteins. When exploring the different Eru types, we observed that the amino acid sequence of the CI repressor differed substantially between Stx phages but there were also homologies which were used group them into eight major Clades (I-VIII). In phage lambda, CI represses expression of upstream genes by forming dimers which bind to specific promoter sequences and self-cleavage of CI relieves the repression (15, 20). All CI proteins belonging to Clade I-VII exhibit the self-catalytic Peptidase_S24 domain and the lambda S150 autocleavage residue (16, 31, 36) which mediates the cleavage of CI resulting in relieve of repression of the promoters in the replication module. However, CI sequences belonging to Clade VIIIa lack this domain and it remains unexplored how this atypical CI protein is involved in regulating phage replication. Another atypical CI protein, lacking the HTH DNA-binding domain, was observed in Clade III, and the regulatory functions of this protein is also unknown. Considering the likelihood that CI is involved in regulation of upstream genes, the differences in amino acid sequence observed between CI repressors of different Eru types may reflect adaptation of binding specificities to match distinct target sequences. It is also likely that the differences observed between CI repressors may influence their regulatory network which, in turn, may influence the stability of the lysogenic state and the pathogenic potential of the host EHEC strain.

Stx phages are known to be highly mosaic and composed of gene segments with different evolutionary histories acquired through a variety of mechanisms, such as homologous recombination, transposition, and site-specific recombination (37-39). The variation in CI protein sequence and Eru types and the different combinations of these revealed in the present study, indicates that Stx phages continuously change and that their classification may be less restricted to specific serotypes than previously anticipated (10). We have previously suggested that the Eru2 type may be restricted to serotype O157:H7 and is predominant for the less potent subtype Stx2c phages (10). However, we observed that among the 63 Eru2 phages detected in this study, fourteen were carried by *E. coli* of serotype O157:H7, while the remaining 49 phages (48 in Japanese EHEC strains (Table S1 in supplemental material) and one in a Dutch EHEC strain (PRNJA285020 strains STEC 564; Table S2 in supplemental material)) were carried by *E. coli* of serotype O121:H19. All Eru2 phages carried by O121:H19 strains encoded Stx2a, while all the O157:H7 strains carried Eru2 phages encoding Stx2c. We also observed that five of the six highly pathogenic strains of serotype O157:H7, which have caused large outbreaks in the UK carry Eru2 phages, and that four of these Eru2 phages encode Stx2c (Table 4). Although, the UK outbreak strains also do carry phages encoding the more potent Stx2a. All in all, this indicate that Eru2 phages are not restricted to hosts of serotype O157:H7 but Eru2 phages carried by this serotype predominantly encode the Stx2c subtype.

Surprisingly, we did not observe any Eru3 type Stx phages among the European STEC strains examined during this study (Table 3). We have previously shown that Eru3 phages were carried by both serotype O157:H7 and O111 strains and often encode the potent subtype Stx2a (10). A majority of the Eru3 type of Stx phages described in the previous work were isolated in the US, indicating that this phage type may be more widespread on the American continent than in Europe.

*E. coli* may carry multiple *stx* negative prophages with similarities to Stx phages together with multiple Stx phages in its genome (40). Therefore, identification of the Eru type requires that the *stx* genes and the phage replication region is present on the same contig or scaffold. Assessment of Eru type from genome sequences generated by short read sequencing technology is often impossible due to contig breaks in the region between *cI* and *stx* (ND in Table S2 in supplemental material). Stx phages often carry repetitive tRNA encoding genes immediately upstream of the *stx* making assembly of contigs difficult in this region.

In the present study, we observe that the Stx2a encoding phages carried by highly virulent EHEC strains from UK (28) and the HUS causing strains from Norway (29) are of Eru1-, Eru2-, Eru5- and Eru7-types. We have previously shown that the Eru1 type is carried by highly pathogenic EHEC strains and that Eru1 phages exhibit a less stable lysogenic state than the classical lambdoid Stx phages (10). It is already well known that the outcome of EHEC disease is often more severe when the infection is caused by an *E. coli* strain producing Stx2 compared to a strain producing Stx1 (3, 5). We must, however, emphasize that the amount of toxin produced must be taken into consideration. It is therefore of great importance to gain more knowledge about how the gene content of the replication region influences regulation of the phage life cycle and, consequently, the levels of Stx produced. More research is also needed to understand how different CI repressor types react to environmental stressors such as the host immune system and antibiotic treatment and the impact of these factors on the Stx production. Importantly, this work highlights that our understanding of bacterial pathogens cannot solely be based on studies on a few model bacterial strains and/or phage types.

## MATERIAL AND METHODS

A total of 120 Stx-converting phage genome sequences were retrieved from the NCBI virus database (taxid:10239) by Standard Nucleotide BLAST using the A subunit of *stx1* (M19437.1) and *stx2* (AF125520) as query sequences (August 2021) (Table S1 in supplemental material).

In addition, ten different bio-projects comprising European STEC strains, one Dutch (PRJNA285020), one Norwegian (PRJEB6447), one French (PRPRJNA706995), three Swiss (PRJNA680568, PRJNA694525, PRJNA438214), one English (PRJNA248042), one Italian (PRJNA666781), one German (PRJNA715185) and one Portuguese (PRJNA643688), were examined for contigs containing *stx* using BLAST as described above (Table S2 in supplemental material). The dataset contained more than 3000 STEC isolates, however, the majority of contigs were too short (<8000 bp) to contain *cI* and *stx* genes on the same contig thus only contigs larger than 8000 bp were examined. A total of 299 contigs containing the region between the CI-coding gene and the *stx* genes were identified in the dataset. The sequences were examined using pDRAW and Eru types were defined by the proteins encoded by the two genes located directly upstream of *cI*. GenomeNet Motif Search (Kyoto University Bioinformatics Center) was used for detection of protein motifs (41). Erus were numbered consecutively as they were detected.

The 260 CI protein sequences (Table S3 in supplemental material), mainly extracted from the abovementioned nucleotide seqcuences, were aligned using ClustalOmega (42). A maximum likelihood tree was inferred from the alignment using IQ-TREE v1.6.12 (43). Node supports were evaluated using the option -bb for ultrafast bootstraps (44) and the VT+GT model was selected as the best evolutionary model using ModelFinder and the BIC criterion (45). Interactive Tree Of Life (iTOL) v6.4 was used for visualization (46).

## List of supplemental information

Table S1: Stx-converting phage genomes with Eru type

Table S2: BioProjects comprising European STEC strains with Eru type

Table S3: Information about the 260 CI sequences used in phylogenetic analysis

Table S4: Percent Identity Matrix (Clustal2.1) between clades of CI repressor sequences

## REFERENCES

1. Bielaszewska M, Mellmann A, Zhang W, Köck R, Fruth A, Bauwens A, Peters G, Karch H. 2011. Characterisation of the Escherichia coli strain associated with an outbreak of haemolytic uraemic syndrome in Germany, 2011: a microbiological study. Lancet Infect Dis 11:671–6.

2. Rasko DA, Webster DR, Sahl JW, Bashir A, Boisen N, Scheutz F, Paxinos EE, Sebra R, Chin CS, Iliopoulos D, Klammer A, Peluso P, Lee L, Kislyuk AO, Bullard J, Kasarskis A, Wang S, Eid J, Rank D, Redman JC, Steyert SR, Frimodt-Møller J, Struve C, Petersen AM, Krogfelt KA, Nataro JP, Schadt EE, Waldor MK. 2011. Origins of the E. coli strain causing an outbreak of hemolytic-uremic syndrome in Germany. N Engl J Med 365:709–17.

3. Scheutz F, Teel LD, Beutin L, Piérard D, Buvens G, Karch H, Mellmann A, Caprioli A, Tozzoli R, Morabito S, Strockbine NA, Melton-Celsa AR, Sanchez M, Persson S, O’Brien AD. 2012. Multicenter evaluation of a sequence-based protocol for subtyping Shiga toxins and standardizing Stx nomenclature. J Clin Microbiol 50:2951–63.

4. Boerlin P, McEwen SA, Boerlin-Petzold F, Wilson JB, Johnson RP, Gyles CL. 1999. Associations between virulence factors of Shiga toxin-producing Escherichia coli and disease in humans. J Clin Microbiol 37:497–503.

5. Fuller CA, Pellino CA, Flagler MJ, Strasser JE, Weiss AA. 2011. Shiga toxin subtypes display dramatic differences in potency. Infect Immun 79:1329–37.

6. Zeng L, Skinner SO, Zong C, Sippy J, Feiss M, Golding I. 2010. Decision making at a subcellular level determines the outcome of bacteriophage infection. Cell 141:682–91.

7. Wagner PL, Acheson DW, Waldor MK. 1999. Isogenic lysogens of diverse shiga toxin 2-encoding bacteriophages produce markedly different amounts of shiga toxin. Infect Immun 67:6710–4.

8. O’Brien AD, Marques LR, Kerry CF, Newland JW, Holmes RK. 1989. Shiga-like toxin converting phage of enterohemorrhagic Escherichia coli strain 933. Microb Pathog 6:381–90.

9. Plunkett G, 3rd, Rose DJ, Durfee TJ, Blattner FR. 1999. Sequence of Shiga toxin 2 phage 933W from Escherichia coli O157:H7: Shiga toxin as a phage late-gene product. J Bacteriol 181:1767–78.

10. Llarena AK, Aspholm M, O’Sullivan K, Wêgrzyn G, Lindbäck T. 2021. Replication Region Analysis Reveals Non-lambdoid Shiga Toxin Converting Bacteriophages. Front Microbiol 12:640945.

11. Pinto G, Sampaio M, Dias O, Almeida C, Azeredo J, Oliveira H. 2021. Insights into the genome architecture and evolution of Shiga toxin encoding bacteriophages of Escherichia coli. BMC Genomics 22:366.

12. Bednarz M, Halliday JA, Herman C, Golding I. 2014. Revisiting bistability in the lysis/lysogeny circuit of bacteriophage lambda. PLoS One 9:e100876.

13. Casjens SR, Hendrix RW. 2015. Bacteriophage lambda: Early pioneer and still relevant. Virology 479-480:310–30.

14. LeBowitz JH, Zylicz M, Georgopoulos C, McMacken R. 1985. Initiation of DNA replication on single-stranded DNA templates catalyzed by purified replication proteins of bacteriophage lambda and Escherichia coli. Proc Natl Acad Sci U S A 82:3988–92.

15. Stayrook S, Jaru-Ampornpan P, Ni J, Hochschild A, Lewis M. 2008. Crystal structure of the lambda repressor and a model for pairwise cooperative operator binding. Nature 452:1022–5.

16. Bell CE, Frescura P, Hochschild A, Lewis M. 2000. Crystal structure of the lambda repressor C-terminal domain provides a model for cooperative operator binding. Cell 101:801–11.

17. Pabo CO, Sauer RT, Sturtevant JM, Ptashne M. 1979. The lambda repressor contains two domains. Proc Natl Acad Sci U S A 76:1608–12.

18. Nyíri K, Kőhegyi B, Micsonai A, Kardos J, Vertessy BG. 2015. Evidence-Based Structural Model of the Staphylococcal Repressor Protein: Separation of Functions into Different Domains. PLoS One 10:e0139086.

19. Pabo CO, Lewis M. 1982. The operator-binding domain of lambda repressor: structure and DNA recognition. Nature 298:443–7.

20. Sauer RT, Ross MJ, Ptashne M. 1982. Cleavage of the lambda and P22 repressors by recA protein. J Biol Chem 257:4458–62.

21. Henthorn KS, Friedman DI. 1995. Identification of related genes in phages phi 80 and P22 whose products are inhibitory for phage growth in Escherichia coli IHF mutants. J Bacteriol 177:3185–90.

22. Iyer LM, Koonin EV, Aravind L. 2002. Extensive domain shuffling in transcription regulators of DNA viruses and implications for the origin of fungal APSES transcription factors. Genome Biol 3:Research0012.

23. Núñez B, De La Cruz F. 2001. Two atypical mobilization proteins are involved in plasmid CloDF13 relaxation. Mol Microbiol 39:1088–99.

24. Zou X, Caufield PW, Li Y, Qi F. 2001. Complete nucleotide sequence and characterization of pUA140, a cryptic plasmid from Streptococcus mutans. Plasmid 46:77–85.

25. Lipps G, Weinzierl AO, von Scheven G, Buchen C, Cramer P. 2004. Structure of a bifunctional DNA primase-polymerase. Nat Struct Mol Biol 11:157–62.

26. Goliger JA, Roberts JW. 1987. Bacteriophage 82 gene Q and Q protein. Sequence, overproduction, and activity as a transcription antiterminator in vitro. J Biol Chem 262:11721–5.

27. Grayhack EJ, Roberts JW. 1982. The phage lambda Q gene product: activity of a transcription antiterminator in vitro. Cell 30:637–48.

28. Yara DA, Greig DR, Gally DL, Dallman TJ, Jenkins C. 2020. Comparison of Shiga toxin-encoding bacteriophages in highly pathogenic strains of Shiga toxin-producing Escherichia coli O157:H7 in the UK. Microb Genom 6.

29. Haugum K, Johansen J, Gabrielsen C, Brandal LT, Bergh K, Ussery DW, Drabløs F, Afset JE. 2014. Comparative genomics to delineate pathogenic potential in non-O157 Shiga toxin-producing Escherichia coli (STEC) from patients with and without haemolytic uremic syndrome (HUS) in Norway. PLoS One 9:e111788.

30. Durante-Rodríguez G, Mancheño JM, Díaz E, Carmona M. 2016. Refactoring the λ phage lytic/lysogenic decision with a synthetic regulator. Microbiologyopen 5:575–81.

31. Kim B, Little JW. 1993. LexA and lambda Cl repressors as enzymes: specific cleavage in an intermolecular reaction. Cell 73:1165–73.

32. Mo CY, Birdwell LD, Kohli RM. 2014. Specificity determinants for autoproteolysis of LexA, a key regulator of bacterial SOS mutagenesis. Biochemistry 53:3158–68.

33. Kozłowska K, Glinkowska M, Boss L, Gaffke L, Deptuła J, Węgrzyn G. 2020. Formation of Complexes Between O Proteins and Replication Origin Regions of Shiga Toxin-Converting Bacteriophages. Front Mol Biosci 7:207.

34. Bochow S, Elliman J, Owens L. 2012. Bacteriophage adenine methyltransferase: a life cycle regulator? Modelled using Vibrio harveyi myovirus like. J Appl Microbiol 113:1001–13.

35. Casjens S, Eppler K, Parr R, Poteete AR. 1989. Nucleotide sequence of the bacteriophage P22 gene 19 to 3 region: identification of a new gene required for lysis. Virology 171:588–98.

36. Luo Y, Pfuetzner RA, Mosimann S, Paetzel M, Frey EA, Cherney M, Kim B, Little JW, Strynadka NC. 2001. Crystal structure of LexA: a conformational switch for regulation of self-cleavage. Cell 106:585–94.

37. Hatfull GF, Hendrix RW. 2011. Bacteriophages and their genomes. Curr Opin Virol 1:298–303.

38. Johansen BK, Wasteson Y, Granum PE, Brynestad S. 2001. Mosaic structure of Shiga-toxin-2-encoding phages isolated from Escherichia coli O157:H7 indicates frequent gene exchange between lambdoid phage genomes. Microbiology (Reading) 147:1929–1936.

39. Smith DL, Rooks DJ, Fogg PC, Darby AC, Thomson NR, McCarthy AJ, Allison HE. 2012. Comparative genomics of Shiga toxin encoding bacteriophages. BMC Genomics 13:311.

40. Zhang Y, Liao YT, Salvador A, Sun X, Wu VCH. 2019. Prediction, Diversity, and Genomic Analysis of Temperate Phages Induced From Shiga Toxin-Producing Escherichia coli Strains. Front Microbiol 10:3093.

41. Bateman A, Birney E, Cerruti L, Durbin R, Etwiller L, Eddy SR, Griffiths-Jones S, Howe KL, Marshall M, Sonnhammer EL. 2002. The Pfam protein families database. Nucleic Acids Res 30:276–80.

42. Sievers F, Wilm A, Dineen D, Gibson TJ, Karplus K, Li W, Lopez R, McWilliam H, Remmert M, Söding J, Thompson JD, Higgins DG. 2011. Fast, scalable generation of high-quality protein multiple sequence alignments using Clustal Omega. Mol Syst Biol 7:539.

43. Nguyen LT, Schmidt HA, von Haeseler A, Minh BQ. 2015. IQ-TREE: a fast and effective stochastic algorithm for estimating maximum-likelihood phylogenies. Mol Biol Evol 32:268–74.

44. Hoang DT, Chernomor O, von Haeseler A, Minh BQ, Vinh LS. 2018. UFBoot2: Improving the Ultrafast Bootstrap Approximation. Mol Biol Evol 35:518–522.

45. Kalyaanamoorthy S, Minh BQ, Wong TKF, von Haeseler A, Jermiin LS. 2017. ModelFinder: fast model selection for accurate phylogenetic estimates. Nat Methods 14:587–589.

46. Letunic I, Bork P. 2021. Interactive Tree Of Life (iTOL) v5: an online tool for phylogenetic tree display and annotation. Nucleic Acids Res 49:W293–w296.

47. Tóth I, Sváb D, Bálint B, Brown-Jaque M, Maróti G. 2016. Comparative analysis of the Shiga toxin converting bacteriophage first detected in Shigella sonnei. Infect Genet Evol 37:150–7.

48. Scotland SM, Willshaw GA, Smith HR, Rowe B. 1987. Properties of strains of Escherichia coli belonging to serogroup O157 with special reference to production of Vero cytotoxins VT1 and VT2. Epidemiol Infect 99:613–24.

49. Taylor CM, White RH, Winterborn MH, Rowe B. 1986. Haemolytic-uraemic syndrome: clinical experience of an outbreak in the West Midlands. Br Med J (Clin Res Ed) 292:1513–6.

50. Byrne L, Dallman TJ, Adams N, Mikhail AFW, McCarthy N, Jenkins C. 2018. Highly Pathogenic Clone of Shiga Toxin-Producing Escherichia coli O157:H7, England and Wales. Emerg Infect Dis 24:2303–2308.

51. Gobin M, Hawker J, Cleary P, Inns T, Gardiner D, Mikhail A, McCormick J, Elson R, Ready D, Dallman T, Roddick I, Hall I, Willis C, Crook P, Godbole G, Tubin-Delic D, Oliver I. 2018. National outbreak of Shiga toxin-producing Escherichia coli O157:H7 linked to mixed salad leaves, United Kingdom, 2016. Euro Surveill 23.

