## Supplemental Tables for "High diversity in the regulatory region of Stx-converting bacteriophage genomes"

### Supplemental Table S1

#### Stx-converting phage genomes with Eru type

| Phage | Serotype | Acc no | Country | Year | Stx type | Eru type |
|---|---|---|---|---|---|---|
| Shigella phage 75/02 Stx | Shigella | NC_029120.1 | Ungarn | 2013 | Stx1 | Eru4 |
| Shigella phage POCJ13 | Shigella | KJ603229 | USA | 2014 | Stx1 | Eru4 |
| Stx2 Converting phage I | O157:H7 | AP004402 | Japan | 2001 | Stx2a | lambdoid |
| Stx2 Converting phage II | O157:H7 | AP005154 | Japan | 2002 | Stx2 | Eru3 |
| Stx2 Converting phage 1717 | O157:H7 | FJ188381 | Canada | 2008 | Stx2a | Eru2 |
| Escherichia phage PA4 | O157:H7 | KP682372.1 | USA | 2015 | Stx2a | Eru3 |
| Escherichia phage PA5 | O157:H7 | KP682373.1 | USA | 2015 | Stx2a | Eru3 |
| Escherichia phage PA11 | O157:H7 | KP682375.1 | USA | 2015 | Stx2a | Eru3 |
| Escherichia phage PA12 | O157:H7 | KP682376.1 | USA | 2015 | Stx2a | Eru3 |
| Escherichia phage PA16 | O157:H7 | KP682377.1 | USA | 2015 | Stx2a | Eru3 |
| Escherichia phage PA21 | O157:H7 | KP682379.1 | USA | 2015 | Stx2a | Eru3 |
| Escherichia phage PA27 | O157:H7 | KP682380.1 | USA | 2015 | Stx2a | Eru3 |
| Escherichia phage PA28 | O157:H7 | KP682381.1 | USA | 2015 | Stx2a | lambdoid |
| Escherichia phage PA29 | O157:H7 | KP682382.1 | USA | 2015 | Stx2a | Eru3 |
| Escherichia phage PA36 | O157:H7 | KP682386.1 | USA | 2015 | Stx2a | Eru3 |
| Escherichia phage PA42 | O157:H7 | KP682387.1 | USA | 2015 | Stx2a | Eru3 |
| Escherichia phage PA45 | O157:H7 | KP682389.1 | USA | 2015 | Stx2a | Eru3 |
| Escherichia phage PA50 | O157:H7 | KP682390.1 | USA | 2015 | Stx2a | Eru3 |
| Escherichia phage PA52 | O157:H7 | KP682392.1 | USA | 2015 | Stx2a | Eru3 |
| Escherichia phage P13771 | O104:H4 | HG792104.1 | Germany | 2009 | Stx2a | Eru1 |
| Escherichia phage P14437 | O104:H4 | HG792105.1 | Norway | 2006 | Stx2a | Eru1 |
| Escherichia phage P13363 | O104:H4 | HG803182.1 | Germany | 2011 | Stx2a | Eru1 |
| Escherichia phage P13374 proviral | O104:H4 | HE664024.1 | Germany | 2011 | Stx2a | Eru1 |
| Enterobacteria phage YYZ-2008 | O157:H7 | FJ184280 | Canada | 2008 | Stx1 | Eru2 |
| Escherichia phage SH2026Stx1 | O157:H7 | NC_049919.1 | USA | 2018 | Stx1 | Eru2 |
| Escherichia phage GER2 | O117:H7 | MG710528.1 | UK | 2017 | Stx1 | Eru1 |
| Escherichia Stx1-converting recombinant phage HUN/2013 | O157:H7 | KJ909655.1 | Hungary | 2013 | Stx1 | Eru3 |
| Stx1 converting phage | O157:H7 | AP005153.1 | Japan | 2003 | Stx1 | Eru3 |
| Bacteriophage CP-1639 and chromosomal integration site | O111:H- | AJ304858.2 | Germany | 2000 | Stx1 | Eru3 |
| Phage BP-4795 complete genome | O84:H4 | AJ556162.1 | Germany | 2003 | Stx1 | lambdoid |
| Stx2-converting phage Stx2a_WGPS9 proviral | O157:H7 | AP012535.1 | Japan | 2012 | Stx2a | lambdoid |
| Stx2-converting phage Stx2a_F403 proviral | O157:H7 | AP012529.1 | Japan | 2012 | Stx2a | Eru5 |
| Stx2-converting phage Stx2a_F422 proviral | O157:H7 | AP012531.1 | Japan | 2012 | Stx2a | lambdoid |
| Stx2-converting phage Stx2a_F451 proviral | O157:H7 | AP012532.1 | Japan | 2012 | Stx2a | Eru5 |
| Stx2-converting phage Stx2a_F723 proviral | O157:H7 | AP012533.1 | Japan | 2012 | Stx2a | lambdoid |
| Escherichia phage P13803 | O2:H27 | HG792102.1 | Germany | 2013 | Stx2a | Eru1 |

| | | | | | | |
|---|---|---|---|---|---|---|
| Stx2-converting phage Stx2a_F765 proviral | O157:H7 | AP012534.1 | Japan | 2012 | Stx2a | Eru1 |
| Stx2-converting phage 86 | O86:H- | NC_008464.1 | USA | 2003 | Stx2a | Eru3 |
| Stx2-converting phage Stx2a_WGPS8 proviral | O157:H7 | AP012540.1 | Japan | 2012 | Stx2c | Eru2 |
| Stx2-converting phage Stx2a_WGPS4 proviral | O157:H7 | AP012538.1 | Japan | 2012 | Stx2c | Eru2 |
| Stx2-converting phage Stx2a_F349 proviral | O157:H7 | AP012530.1 | Japan | 2012 | Stx2c | Eru2 |
| Stx2-converting phage 1717, complete prophage genome | O157:H7 | NC_011357.1 | Canada | 2008 | Stx2c | Eru2 |
| Enterobacteria phage 2851 | O157:H7 | FM180578.1 | Germany | 1993 | Stx2c | Eru2 |
| Stx2-converting phage Stx2a_WGPS6 proviral | O157:H7 | AP012539.1 | Japan | 2012 | Stx2c | Eru2 |
| Stx2-converting phage Stx2a_WGPS2 proviral | O157:H7 | AP012537.1 | Japan | 2012 | Stx2c | Eru6 |
| Stx2-converting phage Stx2a_1447 proviral | O157:H7 | AP012536.1 | Japan | 2012 | Stx2c | Eru6 |
| Stx2 converting phage vB_EcoP_24B | O157:H7 | HM208303.1 | UK | 2010 | Stx2 | Eru5 |
| Enterobacteria phage Min27 | O157:H7 | NC_010237.1 | China | 2007 | Stx2a | Eru5 |
| Lys8385Vzw | O103:H11 | MT225100 | Japan | 2020 | Stx1 | Eru6 |
| Lys19259Vzw | O157:H7 | MT225101 | Japan | 2020 | Stx2 | lambdoid |
| Bacteriophage P27 | Ont:H- | AJ298298.1 | Germany | 2001 | stx2e | Eru7 |
| Stx1 converting phage AU5Stx1 | O157 | KU977419.1 | Australia | 2016 | Stx1 | lambdoid |
| Stx1 converting phage AU6Stx1 | O157 | KU977420.1 | Australia | 2016 | Stx1 | lambdoid |
| Stx2a-converting phage Stx2_14040 | O145:H28 | LC567818.1 | Japan | 2020 | Stx2a | Eru7 |
| Stx1a-converting phage Stx1_14040 | O145:H28 | LC567819.1 | Japan | 2020 | Stx1 | Eru7 |
| Stx2a-converting phage Stx2_14744 | O145:H28 | LC567820.1 | Japan | 2020 | Stx2a | Eru7 |
| Stx1a-converting phage Stx1_14744 | O145:H28 | LC567821.1 | Japan | 2020 | Stx1 | Eru7 |
| Stx1a-converting phage Stx1_132418 | O145:H28 | LC567823.1 | Japan | 2020 | Stx1 | Eru7 |
| Stx2a-converting phage Stx2_499 | O145:H28 | LC567824.1 | Japan | 2020 | Stx2a | Eru7 |
| Stx1a-converting phage Stx1_499 | O145:H28 | LC567825.1 | Japan | 2020 | Stx1 | Eru7 |
| Stx1a-converting phage Stx1_699 | O145:H28 | LC567827.1 | Japan | 2020 | Stx1 | Eru7 |
| Stx2a-converting phage Stx2_EH2201 | O145:H28 | LC567829.1 | Japan | 2020 | Stx2a | Eru7 |
| Stx2d-converting phage Stx2_112808 | O145:H28 | LC567830.1 | Japan | 2020 | Stx2a | Eru7 |
| Stx1a-converting phage Stx1_EH1995 | O145:H28 | LC567831.1 | Japan | 2020 | Stx1 | Eru7 |
| Stx1a-converting phage Stx1_EH1992 | O145:H28 | LC567833.1 | Japan | 2020 | Stx1 | Eru7 |
| Stx2a-converting phage Stx2_EH1910 | O145:H28 | LC567834.1 | Japan | 2020 | Stx2a | Eru7 |
| Stx2a-converting phage Stx2_EH2246 | O145:H28 | LC567837.1 | Japan | 2020 | Stx2a | Eru7 |
| Stx2a-converting phage Stx2_95 | O145:H28 | LC567838.1 | Japan | 2020 | Stx2a | Eru7 |
| Stx2a-converting phage Stx2_EH1846 | O145:H28 | LC567840.1 | Japan | 2020 | Stx2a | Eru7 |
| Stx2a-converting phage Stx2_12E129_PPompW | O145:H28 | LC567841.1 | Japan | 2020 | Stx2a | Eru7 |
| Stx2a-converting phage Stx2_12E129_yecE | O145:H28 | LC567842.1 | Japan | 2020 | Stx2a | Eru7 |
| Stx2a-converting phage Stx2_EH0505 | O121:H19 | LC616031.1 | Japan | 2021 | Stx2a | Eru2 |
| Stx2a-converting phage Stx2_EH1965 | O121:H19 | LC616032.1 | Japan | 2021 | Stx2a | Eru2 |
| Stx2a-converting phage Stx2_EH0787 | O121:H19 | LC616033.1 | Japan | 2021 | Stx2a | Eru2 |
| Stx2a-converting phage Stx2_EH0337 | O121:H19 | LC616034.1 | Japan | 2021 | Stx2a | Eru2 |
| Stx2a-converting phage Stx2_13E027 | O121:H19 | LC616035.1 | Japan | 2021 | Stx2a | Eru2 |
| Stx2a-converting phage Stx2_PV06-102 | O121:H19 | LC616036.1 | Japan | 2021 | Stx2a | Eru2 |

| | | | | | | |
|---|---|---|---|---|---|---|
| Stx2a-converting phage Stx2_4151 | O121:H19 | LC616037.1 | Japan | 2021 | Stx2a | Eru2 |
| Stx2a-converting phage Stx2_707 | O121:H19 | LC616038.1 | Japan | 2021 | Stx2a | Eru2 |
| Stx2a-converting phage Stx2_131033 | O121:H19 | LC616039.1 | Japan | 2021 | Stx2a | Eru2 |
| Stx2a-converting phage Stx2_3993 | O121:H19 | LC616040.1 | Japan | 2021 | Stx2a | Eru2 |
| Stx2a-converting phage Stx2_PV12-16 | O121:H19 | LC616041.1 | Japan | 2021 | Stx2a | Eru2 |
| Stx2a-converting phage Stx2_12E064 | O121:H19 | LC616042.1 | Japan | 2021 | Stx2a | Eru2 |
| Stx2a-converting phage Stx2_3725 | O121:H19 | LC616043.1 | Japan | 2021 | Stx2a | Eru2 |
| Stx2a-converting phage Stx2_8243 | O121:H19 | LC616044.1 | Japan | 2021 | Stx2a | Eru2 |
| Stx2a-converting phage Stx2_481 | O121:H19 | LC616045.1 | Japan | 2021 | Stx2a | Eru2 |
| Stx2a-converting phage Stx2_5122 | O121:H19 | LC616046.1 | Japan | 2021 | Stx2a | Eru2 |
| Stx2a-converting phage Stx2_6804 | O121:H19 | LC616047.1 | Japan | 2021 | Stx2a | Eru2 |
| Stx2a-converting phage Stx2_07Y06 | O121:H19 | LC616048.1 | Japan | 2021 | Stx2a | Eru2 |
| Stx2a-converting phage Stx2_11Y11 | O121:H19 | LC616049.1 | Japan | 2021 | Stx2a | Eru2 |
| Stx2a-converting phage Stx2_3350 | O121:H19 | LC616050.1 | Japan | 2021 | Stx2a | Eru2 |
| Stx2a-converting phage Stx2_PV07-173 | O121:H19 | LC616051.1 | Japan | 2021 | Stx2a | Eru2 |
| Stx2a-converting phage Stx2_7982 | O121:H19 | LC616052.1 | Japan | 2021 | Stx2a | Eru2 |
| Stx2a-converting phage Stx2_12849 | O121:H19 | LC616053.1 | Japan | 2021 | Stx2a | Eru2 |
| Stx2a-converting phage Stx2_12E092 | O121:H19 | LC616054.1 | Japan | 2021 | Stx2a | Eru2 |
| Stx2a-converting phage Stx2_1603 | O121:H19 | LC616055.1 | Japan | 2021 | Stx2a | Eru2 |
| Stx2a-converting phage Stx2_3772 | O121:H19 | LC616056.1 | Japan | 2021 | Stx2a | Eru2 |
| Stx2a-converting phage Stx2_579 | O121:H19 | LC616057.1 | Japan | 2021 | Stx2a | Eru2 |
| Stx2a-converting phage Stx2_716 | O121:H19 | LC616058.1 | Japan | 2021 | Stx2a | Eru2 |
| Stx2a-converting phage Stx2_3105 | O121:H19 | LC616059.1 | Japan | 2021 | Stx2a | Eru2 |
| Stx2a-converting phage Stx2_3417 | O121:H19 | LC616060.1 | Japan | 2021 | Stx2a | Eru2 |
| Stx2a-converting phage Stx2_12874 | O121:H19 | LC616061.1 | Japan | 2021 | Stx2a | Eru2 |
| Stx2a-converting phage Stx2_3896 | O121:H19 | LC616062.1 | Japan | 2021 | Stx2a | Eru2 |
| Stx2a-converting phage Stx2_3418 | O121:H19 | LC616063.1 | Japan | 2021 | Stx2a | Eru2 |
| Stx2a-converting phage Stx2_453 | O121:H19 | LC616064.1 | Japan | 2021 | Stx2a | Eru2 |
| Stx2a-converting phage Stx2_463 | O121:H19 | LC616065.1 | Japan | 2021 | Stx2a | Eru2 |
| Stx2a-converting phage Stx2_11106 | O121:H19 | LC616066.1 | Japan | 2021 | Stx2a | Eru2 |
| Stx2a-converting phage Stx2_356 | O121:H19 | LC616067.1 | Japan | 2021 | Stx2a | Eru2 |
| Stx2a-converting phage Stx2_522 | O121:H19 | LC616068.1 | Japan | 2021 | Stx2a | Eru2 |
| Stx2a-converting phage Stx2_635 | O121:H19 | LC616069.1 | Japan | 2021 | Stx2a | Eru2 |
| Stx2a-converting phage Stx2_15436 | O121:H19 | LC616070.1 | Japan | 2021 | Stx2a | Eru2 |
| Stx2a-converting phage Stx2_8048 | O121:H19 | LC616071.1 | Japan | 2021 | Stx2a | Eru2 |
| Stx2a-converting phage Stx2_PV03-14 | O121:H19 | LC616072.1 | Japan | 2021 | Stx2a | Eru2 |
| Stx2a-converting phage Stx2_08E027 | O121:H19 | LC616073.1 | Japan | 2021 | Stx2a | Eru2 |
| Stx2a-converting phage Stx2_10902 | O121:H19 | LC616074.1 | Japan | 2021 | Stx2a | Eru2 |
| Stx2a-converting phage Stx2_06E050 | O121:H19 | LC616075.1 | Japan | 2021 | Stx2a | Eru2 |
| Stx2a-converting phage Stx2_2935 | O121:H19 | LC616076.1 | Japan | 2021 | Stx2a | Eru2 |
| Stx2a-converting phage Stx2_13518 | O121:H19 | LC616077.1 | Japan | 2021 | Stx2a | Eru2 |

| | | | | | | |
|---|---|---|---|---|---|---|
| Stx2a-converting phage Stx2_10E082 | O121:H19 | LC616078.1 | Japan | 2021 | Stx2a | Eru2 |
| Stx2a-converting phage Stx2_KH16-043 | O121:H19 | LC616079.1 | Japan | 2021 | Stx2a | Eru7 |

### Table S2

BioProjects comprising European STEC strains with Eru type

| **PRJNA285020**<br>**Shiga toxin-producing *Escherichia coli* isolates obtained from two Dutch regions** | | |
|---|---|---|
| **Strain** | **NCBI Accession no** | **Eru type** |
| **Stx2** | | |
| Escherichia coli strain E09/10 contig_97, whole genome shotgun sequence | LGBK01000097.1 | ND |
| Escherichia coli strain STEC 1109 contig_168, whole genome shotgun sequence | LGBC01000168.1 | ND |
| Escherichia coli strain STEC 1255 STEC1255_contig_195, whole genome shotgun sequence | LOGC01000107.1 | ND |
| Escherichia coli strain STEC 1293 STEC1293_contig_49, whole genome shotgun sequence | LOGF01000197.1 | Eru7 |
| Escherichia coli strain STEC 1299 STEC1299_contig_22, whole genome shotgun sequence | LOGG01000057.1 | Eru6 |
| Escherichia coli strain STEC 1363 STEC1363_contig_23, whole genome shotgun sequence | LOGI01000090.1 | Eru6 |
| Escherichia coli strain STEC 1375 STEC1375_contig_25, whole genome shotgun sequence | LOGJ01000018.1 | Eru7 |
| Escherichia coli strain STEC 1442 STEC1442_contig_2, whole genome shotgun sequence | LOGK01000083.1 | Eru7 |
| Escherichia coli strain STEC 1506 STEC1506_contig_10, whole genome shotgun sequence | LPWV01000002.1 | ND |
| Escherichia coli strain STEC 1528 STEC1528_contig_43, whole genome shotgun sequence | LOGP01000038.1 | ND |
| Escherichia coli strain STEC 1634 STEC1634_contig_43, whole genome shotgun sequence | LOGS01000097.1 | ND |
| Escherichia coli strain STEC 1686 STEC1686_contig_77, whole genome shotgun sequence | LOGT01000177.1 | Eru10 |
| Escherichia coli strain STEC 200 STEC-200_contig_39, whole genome shotgun sequence | LNZL01000033.1 | Eru7 |
| Escherichia coli strain STEC 2075 contig_86, whole genome shotgun sequence | LGBD01000086.1 | ND |
| Escherichia coli strain STEC 2110.1 2110-1_contig_200, whole genome shotgun sequence | LPWW01000114.1 | ND |
| Escherichia coli strain STEC 2110.1 2110-1_contig_204, whole genome shotgun sequence | LPWW01000118.1 | ND |
| Escherichia coli strain STEC 2110.1 2110-1_contig_232, whole genome shotgun sequence | LPWW01000149.1 | ND |
| Escherichia coli strain STEC 2110.1 2110-1_contig_40, whole genome shotgun sequence | LPWW01000216.1 | ND |
| Escherichia coli strain STEC 2112 contig_150, whole genome shotgun sequence | LGBE01000150.1 | ND |
| Escherichia coli strain STEC 2236 STEC2236_contig_23, whole genome shotgun sequence | LOGY01000073.1 | lambdoid |
| Escherichia coli strain STEC 2257 contig_109, whole genome shotgun sequence | LGBF01000109.1 | ND |
| Escherichia coli strain STEC 2410 contig_189, whole genome shotgun sequence | LGBG01000189.1 | ND |
| Escherichia coli strain STEC 2410 contig_203, whole genome shotgun sequence | LGBG01000203.1 | ND |
| Escherichia coli strain STEC 2410 contig_72, whole genome shotgun sequence | LGBG01000072.1 | Eru1 |
| Escherichia coli strain STEC 2450 STEC2450_contig_65, whole genome shotgun sequence | LPXA01000177.1 | Eru6 |
| Escherichia coli strain STEC 2499 STEC2499_contig_42, whole genome shotgun sequence | LOIE01000070.1 | ND |
| Escherichia coli strain STEC 2505 STEC2505_contig_24, whole genome shotgun sequence | LPXB01000100.1 | Eru6 |
| Escherichia coli strain STEC 2573 STEC2573_contig_104, whole genome shotgun sequence | LOIH01000007.1 | ND |
| Escherichia coli strain STEC 2573 STEC2573_contig_138, whole genome shotgun sequence | LOIH01000044.1 | ND |
| Escherichia coli strain STEC 2573 STEC2573_contig_62, whole genome shotgun sequence | LOIH01000107.1 | ND |
| Escherichia coli strain STEC 2573 STEC2573_contig_7, whole genome shotgun sequence | LOIH01000115.1 | ND |
| Escherichia coli strain STEC 2591 STEC2591_contig_56, whole genome shotgun sequence | LOII01000068.1 | ND |
| Escherichia coli strain STEC 2595 STEC2595_contig_34, whole genome shotgun sequence | LOIJ01000033.1 | Eru11 |

| | | |
|---|---|---|
| Escherichia coli strain STEC 2620 STEC2620_contig_169, whole genome shotgun sequence | LPXC01000078.1 | ND |
| Escherichia coli strain STEC 2620 STEC2620_contig_226, whole genome shotgun sequence | LPXC01000142.1 | ND |
| Escherichia coli strain STEC 2620 STEC2620_contig_69, whole genome shotgun sequence | LPXC01000201.1 | ND |
| Escherichia coli strain STEC 2667 contig_19, whole genome shotgun sequence | LGBH01000019.1 | ND |
| Escherichia coli strain STEC 2746 STEC2746_contig_61, whole genome shotgun sequence | LPXD01000148.1 | Eru6 |
| Escherichia coli strain STEC 2770 STEC2770_contig_18, whole genome shotgun sequence | LPXF01000088.1 | ND |
| Escherichia coli strain STEC 2797 STEC2797_contig_85, whole genome shotgun sequence | LOIP01000113.1 | ND |
| Escherichia coli strain STEC 2820 contig_161, whole genome shotgun sequence | LGBQ01000161.1 | ND |
| Escherichia coli strain STEC 2821 contig_98, whole genome shotgun sequence | LGBI01000098.1 | ND |
| Escherichia coli strain STEC 2826 STEC2826_contig_97, whole genome shotgun sequence | LOJA01000134.1 | ND |
| Escherichia coli strain STEC 2861 STEC2861_contig_8, whole genome shotgun sequence | LOIR01000058.1 | Eru11 |
| Escherichia coli strain STEC 2868 contig_83, whole genome shotgun sequence | LGBJ01000083.1 | ND |
| Escherichia coli strain STEC 2894.2 2894-2_contig_216, whole genome shotgun sequence | LOIT01000131.1 | ND |
| Escherichia coli strain STEC 2953 STEC2953_contig_5, whole genome shotgun sequence | LOIW01000084.1 | Eru10 |
| Escherichia coli strain STEC 2954 STEC2954_contig_124, whole genome shotgun sequence | LPXE01000029.1 | ND |
| Escherichia coli strain STEC 2980 STEC2980_contig_28, whole genome shotgun sequence | LOIY01000123.1 | Eru6 |
| Escherichia coli strain STEC 3031 STEC3031_contig_25, whole genome shotgun sequence | LOIZ01000129.1 | Eru6 |
| Escherichia coli strain STEC 3039 STEC3039_contig_118, whole genome shotgun sequence | LPUH01000022.1 | Eru6 |
| Escherichia coli strain STEC 3039 STEC3039_contig_29, whole genome shotgun sequence | LPUH01000108.1 | Eru6 |
| Escherichia coli strain STEC 3055 STEC3055_contig_130, whole genome shotgun sequence | LPUI01000036.1 | ND |
| Escherichia coli strain STEC 3055 STEC3055_contig_149, whole genome shotgun sequence | LPUI01000056.1 | ND |
| Escherichia coli strain STEC 3055 STEC3055_contig_201, whole genome shotgun sequence | LPUI01000115.1 | ND |
| Escherichia coli strain STEC 3098 STEC3098_contig_22, whole genome shotgun sequence | LPUM01000015.1 | Eru6 |
| Escherichia coli strain STEC 343 contig_50, whole genome shotgun sequence | LDOZ01000050.1 | ND |
| Escherichia coli strain STEC 440 440_contig_159, whole genome shotgun sequence | MRVY01000159.1 | ND |
| Escherichia coli strain STEC 476-14 476-14_contig_49, whole genome shotgun sequence | MRVU01000049.1 | Eru7 |
| Escherichia coli strain STEC 479BS2 479BS2_contig_11, whole genome shotgun sequence | MRVR01000011.1 | Eru7 |
| Escherichia coli strain STEC 480-3 480-3_contig_132, whole genome shotgun sequence | MRVT01000132.1 | ND |
| Escherichia coli strain STEC 510-5 510-5_contig_40, whole genome shotgun sequence | MRVV01000040.1 | Eru7 |
| Escherichia coli strain STEC 514-2 514-2_contig_122, whole genome shotgun sequence | MRVZ01000122.1 | Eru7 |
| Escherichia coli strain STEC 536-9 Re_536-9_contig_20, whole genome shotgun sequence | MRVS01000020.1 | Eru7 |
| Escherichia coli strain STEC 545 STEC-545_contig_46, whole genome shotgun sequence | LODB01000401.1 | Eru8 |
| Escherichia coli strain STEC 563 STEC-563_contig_138, whole genome shotgun sequence | LODD01000044.1 | ND |
| Escherichia coli strain STEC 563 STEC-563_contig_148, whole genome shotgun sequence | LODD01000055.1 | ND |
| Escherichia coli strain STEC 565 STEC565_contig_23, whole genome shotgun sequence | LODE01000106.1 | Eru2 |
| Escherichia coli strain STEC 573-4 573-4_contig_15, whole genome shotgun sequence | MRWA01000015.1 | Eru7 |
| Escherichia coli strain STEC 587-5 587-5_contig_81, whole genome shotgun sequence | MRWB01000081.1 | ND |
| Escherichia coli strain STEC 587-5 587-5_contig_82, whole genome shotgun sequence | MRWB01000082.1 | ND |
| Escherichia coli strain STEC 605 STEC-605_contig_136, whole genome shotgun sequence | LFUA01000136.1 | ND |
| Escherichia coli strain STEC 621S1 621S1_contig_13, whole genome shotgun sequence | MRVX01000013.1 | ND |
| Escherichia coli strain STEC 623 STEC-623_contig_109, whole genome shotgun sequence | LFUB01000109.1 | ND |

| | | |
|---|---|---|
| Escherichia coli strain STEC 625C-4 625C-4_contig_16, whole genome shotgun sequence | MRVW01000016.1 | Eru7 |
| Escherichia coli strain STEC 66 STEC-66_contig_79, whole genome shotgun sequence | LNFT01000172.1 | Eru1 |
| Escherichia coli strain STEC 709 STEC-709_contig_1, whole genome shotgun sequence | LOFM01000001.1 | Eru1 |
| Escherichia coli strain STEC 709 STEC-709_contig_121, whole genome shotgun sequence | LOFM01000026.1 | Eru7 |
| Escherichia coli strain STEC 731 STEC-731_contig_50, whole genome shotgun sequence | LOFN01000117.1 | Eru7 |
| Escherichia coli strain STEC 771 contig_171, whole genome shotgun sequence | LGAZ01000171.1 | ND |
| Escherichia coli strain STEC 915 STEC-915_contig_74, whole genome shotgun sequence | LFUH01000074.1 | ND |
| Escherichia coli strain STEC 931 STEC-931_contig_69, whole genome shotgun sequence | LOFS01000183.1 | Eru1 |
| Escherichia coli strain STEC 989 contig_159, whole genome shotgun sequence | LGBA01000159.1 | ND |
| Escherichia coli strain STEC 994 contig_120, whole genome shotgun sequence | LGBB01000120.1 | ND |
| **Stx1** | | |
| Escherichia coli strain STEC 1117 STEC1117_contig_150, whole genome shotgun sequence | LOFU01000058.1 | ND |
| Escherichia coli strain STEC 1161 STEC1161_contig_128, whole genome shotgun sequence | LOFV01000033.1 | ND |
| Escherichia coli strain STEC 1188 STEC1188_contig_25, whole genome shotgun sequence | LOFX01000035.1 | Eru4 |
| Escherichia coli strain STEC 1201 STEC1201_contig_80, whole genome shotgun sequence | LOFZ01000160.1 | ND |
| Escherichia coli strain STEC 1225 STEC1225_contig_97, whole genome shotgun sequence | LOGA01000204.1 | ND |
| Escherichia coli strain STEC 1236 STEC1236_contig_228, whole genome shotgun sequence | LOGB01000144.1 | ND |
| Escherichia coli strain STEC 1255 STEC1255_contig_18, whole genome shotgun sequence | LOGC01000090.1 | ND |
| Escherichia coli strain STEC 1284 STEC1284_contig_140, whole genome shotgun sequence | LOGE01000047.1 | ND |
| Escherichia coli strain STEC 1293 STEC1293_contig_188, whole genome shotgun sequence | LOGF01000099.1 | ND |
| Escherichia coli strain STEC 1299 STEC1299_contig_7, whole genome shotgun sequence | LOGG01000109.1 | Eru4 |
| Escherichia coli strain STEC 1465 STEC1465_contig_2, whole genome shotgun sequence | LOGL01000097.1 | Eru1 |
| Escherichia coli strain STEC 1500 STEC1500_contig_18, whole genome shotgun sequence | LOGN01000031.1 | Eru4 |
| Escherichia coli strain STEC 1506 STEC1506_contig_62, whole genome shotgun sequence | LPWV01000121.1 | Eru4 |
| Escherichia coli strain STEC 1513 STEC1513_contig_128, whole genome shotgun sequence | LOGO01000033.1 | ND |
| Escherichia coli strain STEC 1532 STEC1532_contig_89, whole genome shotgun sequence | LOGQ01000160.1 | lambdoid |
| Escherichia coli strain STEC 1585 STEC1585_contig_10, whole genome shotgun sequence | LOGR01000002.1 | Eru4 |
| Escherichia coli strain STEC 168 STEC-168_contig_23, whole genome shotgun sequence | LNFV01000026.1 | Eru4 |
| Escherichia coli strain STEC 1686 STEC1686_contig_68, whole genome shotgun sequence | LOGT01000167.1 | ND |
| Escherichia coli strain STEC 169 STEC169_contig_145, whole genome shotgun sequence | LNZJ01000052.1 | ND |
| Escherichia coli strain STEC 196 STEC196_contig_6, whole genome shotgun sequence | LNZK01000067.1 | Eru4 |
| Escherichia coli strain STEC 2064 STEC2064_contig_11, whole genome shotgun sequence | LOJC01000003.1 | Eru4 |
| Escherichia coli strain STEC 2074 STEC2074_contig_35, whole genome shotgun sequence | LOJD01000082.1 | Eru4 |
| Escherichia coli strain STEC 2112 contig_161, whole genome shotgun sequence | LGBE01000161.1 | ND |
| Escherichia coli strain STEC 2144 STEC2144_contig_132, whole genome shotgun sequence | LOGU01000038.1 | lambdoid |
| Escherichia coli strain STEC 2174 STEC2174_contig_4, whole genome shotgun sequence | LOGV01000046.1 | Eru4 |
| Escherichia coli strain STEC 2193 STEC2193_contig_13, whole genome shotgun sequence | LOGW01000035.1 | lambdoid |
| Escherichia coli strain STEC 2211 STEC2211_contig_137, whole genome shotgun sequence | LOGX01000043.1 | ND |
| Escherichia coli strain STEC 2257 contig_143, whole genome shotgun sequence | LGBF01000143.1 | ND |
| Escherichia coli strain STEC 2270 STEC2270_contig_37, whole genome shotgun sequence | LPWX01000147.1 | ND |
| Escherichia coli strain STEC 2346 STEC2346_contig_159, whole genome shotgun sequence | LOHA01000067.1 | ND |

| | | |
|---|---|---|
| Escherichia coli strain STEC 2359 STEC2359_contig_30, whole genome shotgun sequence | LOHB01000024.1 | ND |
| Escherichia coli strain STEC 2363 STEC2363_contig_20, whole genome shotgun sequence | LPWY01000046.1 | Eru4 |
| Escherichia coli strain STEC 2419 STEC2419_contig_65, whole genome shotgun sequence | LPWZ01000120.1 | lambdoid |
| Escherichia coli strain STEC 2441 STEC2441_contig_165, whole genome shotgun sequence | LOHC01000074.1 | ND |
| Escherichia coli strain STEC 2450 STEC2450_contig_19, whole genome shotgun sequence | LPXA01000101.1 | Eru4 |
| Escherichia coli strain STEC 2499 STEC2499_contig_39, whole genome shotgun sequence | LOIE01000066.1 | Eru7 |
| Escherichia coli strain STEC 2505 STEC2505_contig_22, whole genome shotgun sequence | LPXB01000098.1 | Eru4 |
| Escherichia coli strain STEC 2564 STEC2564_contig_44, whole genome shotgun sequence | LOIG01000176.1 | Eru1 |
| Escherichia coli strain STEC 2573 STEC2573_contig_61, whole genome shotgun sequence | LOIH01000106.1 | ND |
| Escherichia coli strain STEC 2591 STEC2591_contig_23, whole genome shotgun sequence | LOII01000032.1 | Eru7 |
| Escherichia coli strain STEC 2620 STEC2620_contig_99, whole genome shotgun sequence | LPXC01000234.1 | ND |
| Escherichia coli strain STEC 2633 STEC2633_contig_39, whole genome shotgun sequence | LOIK01000066.1 | Eru5 |
| Escherichia coli strain STEC 2667 contig_141, whole genome shotgun sequence | LGBH01000141.1 | ND |
| Escherichia coli strain STEC 2708 STEC2708_contig_138, whole genome shotgun sequence | LOIL01000044.1 | ND |
| Escherichia coli strain STEC 2743 STEC2743_contig_90, whole genome shotgun sequence | LOIM01000198.1 | ND |
| Escherichia coli strain STEC 2746 STEC2746_contig_26, whole genome shotgun sequence | LPXD01000109.1 | Eru4 |
| Escherichia coli strain STEC 2764 STEC2764_contig_63, whole genome shotgun sequence | LOIN01000127.1 | Eru4 |
| Escherichia coli strain STEC 2770 STEC2770_contig_19, whole genome shotgun sequence | LPXF01000089.1 | ND |
| Escherichia coli strain STEC 2788 STEC2788_contig_39, whole genome shotgun sequence | LOIO01000033.1 | Eru4 |
| Escherichia coli strain STEC 2820 contig_76, whole genome shotgun sequence | LGBQ01000076.1 | ND |
| Escherichia coli strain STEC 2821 contig_145, whole genome shotgun sequence | LGBI01000145.1 | ND |
| Escherichia coli strain STEC 2839 STEC2839_contig_22, whole genome shotgun sequence | LOJB01000015.1 | Eru4 |
| Escherichia coli strain STEC 2841 STEC2841_contig_11, whole genome shotgun sequence | LOIQ01000003.1 | Eru4 |
| Escherichia coli strain STEC 2868 contig_84, whole genome shotgun sequence | LGBJ01000084.1 | ND |
| Escherichia coli strain STEC 2894.1 2894-1_contig_38, whole genome shotgun sequence | LOIS01000032.1 | Eru5 |
| Escherichia coli strain STEC 29 STEC-29_contig_6, whole genome shotgun sequence | LNFU01000056.1 | Eru4 |
| Escherichia coli strain STEC 2920 STEC2920_contig_172, whole genome shotgun sequence | LOIU01000082.1 | ND |
| Escherichia coli strain STEC 2938 STEC2938_contig_47, whole genome shotgun sequence | LOIV01000111.1 | ND |
| Escherichia coli strain STEC 2953 STEC2953_contig_44, whole genome shotgun sequence | LOIW01000078.1 | Eru4 |
| Escherichia coli strain STEC 2954 STEC2954_contig_9, whole genome shotgun sequence | LPXE01000152.1 | Eru4 |
| Escherichia coli strain STEC 2962 STEC2962_contig_4, whole genome shotgun sequence | LOIX01000034.1 | Eru4 |
| Escherichia coli strain STEC 2980 STEC2980_contig_92, whole genome shotgun sequence | LOIY01000194.1 | Eru4 |
| Escherichia coli strain STEC 299 STEC299_contig_145, whole genome shotgun sequence | LOCR01000052.1 | ND |
| Escherichia coli strain STEC 3031 STEC3031_contig_24, whole genome shotgun sequence | LOIZ01000128.1 | Eru4 |
| Escherichia coli strain STEC 3039 STEC3039_contig_100, whole genome shotgun sequence | LPUH01000003.1 | ND |
| Escherichia coli strain STEC 3055 STEC3055_contig_65, whole genome shotgun sequence | LPUI01000214.1 | Eru1 |
| Escherichia coli strain STEC 3084 STEC3084_contig_28, whole genome shotgun sequence | LPUJ01000040.1 | Eru4 |
| Escherichia coli strain STEC 3087 STEC3087_contig_27, whole genome shotgun sequence | LPUK01000020.1 | Eru4 |
| Escherichia coli strain STEC 3094 STEC3094_contig_88, whole genome shotgun sequence | LPUL01000162.1 | ND |
| Escherichia coli strain STEC 3106 STEC3106_contig_9, whole genome shotgun sequence | LPUN01000125.1 | Eru4 |
| Escherichia coli strain STEC 329 STEC329_contig_68, whole genome shotgun sequence | LOCT01000103.1 | Eru4 |

| | | |
|---|---|---|
| Escherichia coli strain STEC 370 STEC370_contig_124, whole genome shotgun sequence | LOCU01000029.1 | ND |
| Escherichia coli strain STEC 380 STEC380_contig_200, whole genome shotgun sequence | LOCV01000114.1 | ND |
| Escherichia coli strain STEC 384 STEC384_contig_129, whole genome shotgun sequence | LOCW01000034.1 | ND |
| Escherichia coli strain STEC 440 440_contig_83, whole genome shotgun sequence | MRVY01000083.1 | ND |
| Escherichia coli strain STEC 464 STEC464_contig_70, whole genome shotgun sequence | LOCX01000131.1 | ND |
| Escherichia coli strain STEC 477 STEC477_contig_83, whole genome shotgun sequence | LOCY01000237.1 | ND |
| Escherichia coli strain STEC 479 STEC479_contig_147, whole genome shotgun sequence | LOCZ01000054.1 | ND |
| Escherichia coli strain STEC 487 STEC487_contig_273, whole genome shotgun sequence | LODA01000194.1 | ND |
| Escherichia coli strain STEC 559 STEC559_contig_57, whole genome shotgun sequence | LODC01000145.1 | lambdoid |
| Escherichia coli strain STEC 563 STEC-563_contig_155, whole genome shotgun sequence | LODD01000063.1 | ND |
| Escherichia coli strain STEC 621S1 621S1_contig_12, whole genome shotgun sequence | MRVX01000012.1 | ND |
| Escherichia coli strain STEC 623 STEC-623_contig_46, whole genome shotgun sequence | LFUB01000046.1 | ND |
| Escherichia coli strain STEC 627 STEC627_contig_98, whole genome shotgun sequence | LODF01000184.1 | ND |
| Escherichia coli strain STEC 645 STEC645_contig_5, whole genome shotgun sequence | LODG01000045.1 | Eru4 |
| Escherichia coli strain STEC 66 STEC-66_contig_78, whole genome shotgun sequence | LNFT01000171.1 | ND |
| Escherichia coli strain STEC 690 STEC690_contig_253, whole genome shotgun sequence | LOFJ01000172.1 | ND |
| Escherichia coli strain STEC 691 STEC691_contig_122, whole genome shotgun sequence | LOFK01000027.1 | ND |
| Escherichia coli strain STEC 707 STEC707_contig_88, whole genome shotgun sequence | LOFL01000143.1 | ND |
| Escherichia coli strain STEC 709 STEC-709_contig_147, whole genome shotgun sequence | LOFM01000054.1 | ND |
| Escherichia coli strain STEC 757 STEC757_contig_136, whole genome shotgun sequence | LOFO01000042.1 | ND |
| Escherichia coli strain STEC 764 STEC764_contig_242, whole genome shotgun sequence | LOFP01000160.1 | ND |
| Escherichia coli strain STEC 771 contig_117, whole genome shotgun sequence | LGAZ01000117.1 | ND |
| Escherichia coli strain STEC 886 STEC886_contig_19, whole genome shotgun sequence | LOFR01000041.1 | Eru4 |
| Escherichia coli strain STEC 915 STEC-915_contig_344, whole genome shotgun sequence | LFUH01000344.1 | ND |
| Escherichia coli strain STEC 989 contig_82, whole genome shotgun sequence | LGBA01000082.1 | ND |
| **PRJEB6447**<br>**Norwegian STEC isolates** | | |
| **Strain** | **NCBI Accession no** | **Eru type** |
| **Stx2** | | |
| FHI3 | CCPT01000144.1 | Eru7 |
| FHI4 | LM995944.1 | Eru7 |
| FHI5 | CCQC01000098.1 | ND |
| FHI6 | LM996356.1 | Eru1 |
| FHI7 | LM996408.1 | Eru1 |
| FHI9 | LM997071.1 | Eru1 |
| FHI12 | LM995551.1 | Eru1 |
| FHI19 | CCPR01000384.1 | Eru1 |
| FHI22 | CCPS01000056.1 | ND |
| FHI24 | LM995744.1 | Eru1 |
| FHI25 | LM996297.1 | Eru7 |
| FHI27 | LM995668.1 | Eru1 |

| | | |
|---|---|---|
| FHI28 | LM995766.1 | ND |
| FHI30 | LM995896.1 | Eru10 |
| FHI31 | CCVO01000012.1 | ND |
| FHI32 | CCVP01000063.1 | ND |
| FHI35 | CCQJ01000092.1 | ND |
| FHI36 | CCPV01000170.1 | Eru1 |
| FHI37 | CCPU01000077.1 | ND |
| FHI39 | CCPX01000131.1 | Eru6 |
| FHI41 | CCPY01000042.1 | lambdoid |
| FHI42 | LK999941.1 | ND |
| FHI43 | LM996066.1 | Eru7 |
| FHI48 | LM996025.1 | Eru7 (HUS) |
| FHI49 | CCQB01000137.1 | ND |
| FHI53 | CCRA01000038.1 | Eru6 |
| FHI58 | LM995979.1 | ND (HUS) |
| FHI59 | LK999983.1 | Eru6 |
| FHI62 | LN554923.1 | Eru7 |
| FHI63 | LM996460.1 | Eru7 (HUS) |
| FHI65 | LM996529.1 | Eru6 |
| FHI71 | LM996832.1 | Eru6 |
| FHI79 | LM996682.1 | ND (HUS) |
| FHI8 | LM996720.1 | Eru1 (HUS) |
| FHI81 | CCQV01000126.1 | ND |
| FHI82 | LM996798.1 | Eru7 |
| FHI83 | LM996896.1 | Eru2 (HUS) |
| FHI85 | LM996947.1 | Eru1 |
| FHI86 | CCRC01000097.1 | Eru1 |
| FHI88 | CCQX01000142.1 | ND |
| FHI89 | LM997036.1 | Eru6 |
| FHI92 | LM997161.1 | Eru7 |
| FHI95 | CCRD01000099.1 | Eru7 |
| FHI98 | LM997367.1 | Eru10 |
| FHI99 | LM997254.1 | Eru13 |
| FHI100 | LK985420.1 | Eru8 |
| FHI101 | CCPP01000171.1 | Eru7 |
| FHI102 | LM995495.1 | Eru1 (HUS) |
| StOlav104 | LK931573.1 | Eru7 |
| St. Olav164 | PVRW01000040.1 | Eru7 |
| **Stx1** | | |
| FHI1 | CCPO01000158.1 | ND |

| | | |
|---|---|---|
| FHI5 | CCQC01000109.1 | ND |
| FHI6 | LM996340.1 | ND (HUS) |
| FHI7 | LM996413.1 | Eru4 (HUS) |
| FHI20 | LM995817.1 | ND |
| FHI22 | CCPS01000094.1 | Eru4 |
| FHI23 | NZ_LM995659.1 | Eru12 |
| FHI29 | LM995865.1 | Eru4 |
| FHI30 | LM995905.1 | Eru4 |
| FHI32 | CCVP01000073.1 | Eru12 |
| FHI34 | LM996489.1 | Eru4 |
| FHI37 | CCPU01000123.1 | ND |
| FHI40 | LM996246.1 | ND |
| FHI45 | CCQM01000063.1 | Eru4 |
| FHI46 | CCPZ01000238.1 | ND |
| FHI47 | CCQA01000072.1 | Eru6 |
| FHI49 | CCQB01000089.1 | Eru4 |
| FHI50 | CCQD01000134.1 | Eru6 |
| FHI51 | CCQW01000128.1 | ND |
| FHI52 | CCQH01000128.1 | ND |
| FHI54 | CCQI01000086.1 | ND |
| FHI56 | CCQK01000110.1 | ND |
| FHI60 | CCRB01000131.1 | ND |
| FHI61 | CCQL01000336.1 | ND |
| FHI64 | CCQN01000148.1 | ND |
| FHI65 | LM996514.1 | Eru1 |
| FHI67 | CCQO01000029.1 | ND |
| FHI68 | CCQQ01000131.1 | Eru1 |
| FHI69 | CCQP01000089.1 | Eru4 |
| FHI70 | LM997107.1 | Eru2 |
| FHI72 | LM996870.1 | Eru1 |
| FHI74 | LM996604.1 | Eru5 |
| FHI75 | LM996630.1 | ND |
| FHI76 | CCQT01000162.1 | ND |
| FHI77 | CCQS01000108.1 | ND |
| FHI78 | CCQU01000036.1 | ND |
| FHI81 | CCQV01000151.1 | Eru1 |
| FHI84 | CCVQ01000033.1 | ND |
| FHI85 | LM996969.1 | ND |
| FHI87 | LM996984.1 | ND |
| FHI90 | LM997295.1 | ND |
| FHI92 | LM997172.1 | Eru4 |

| | | |
|---|---|---|
| FHI93 | CCQY01000267.1 | ND |
| FHI96 | CCRE01000139.1 | Eru4 |
| FHI97 | LM997224.1 | lambdoid |
| St. Olav143 | CCVR01000362.1 | ND |
| **PRJNA706995**<br>**Shiga-toxin-producing *Escherichia coli* carriage in asymptomatic French infants** | | |
| **Strain** | **NCBI Accession no** | **Eru type** |
| **Stx2** | | |
| Escherichia coli strain 172124GE NODE_96_259616.007695, whole genome shotgun sequence | JAFMZR010000326.1 | ND |
| Escherichia coli strain 192068CE NODE_18_9463935.066618, whole genome shotgun sequence | JAFMZX010000018.1 | Eru7 |
| Escherichia coli strain 182303EI NODE_70377111.543908, whole genome shotgun sequence | JAFMZU010000233.1 | ND |
| Escherichia coli strain 172332DM NODE_73_403113.378842, whole genome shotgun sequence | JAFMZS010000271.1 | ND |
| Escherichia coli strain 182769NA NODE_67_1551313.870207, whole genome shotgun sequence | JAFMZW010000381.1 | Eru7 |
| Escherichia coli strain 172669GA NODE_53_1898931.577163, whole genome shotgun sequence | JAFMZT010000053.1 | Eru6 |
| **Stx1** | | |
| Escherichia coli strain 192068CE NODE_23_7894751.250875, whole genome shotgun sequence | JAFMZX010000023.1 | lambdoid |
| Escherichia coli strain 192161DA NODE_83_706311.409746, whole genome shotgun sequence | JAFMZY010000402.1 | ND |
| Escherichia coli strain 182629BE NODE_112344714.641566, whole genome shotgun sequence | JAFMZV010000014.1 | ND |
| **PRJNA694525**<br>**STEC isolated from raw meat-based diets for companion animals in Switzerland** | | |
| **Strain** | **NCBI Accession no** | **Eru type** |
| **Stx2** | | |
| Escherichia coli strain ATB6-118 contig00027, whole genome shotgun sequence | JAEUYO010000027.1 | ND |
| Escherichia coli strain LSC6-3 contig00094, whole genome shotgun sequence | JAETZC010000094.1 | ND |
| Escherichia coli strain ATC45-11 contig00118, whole genome shotgun sequence | JAETYS010000118.1 | ND |
| Escherichia coli strain ATC46-2 contig00088, whole genome shotgun sequence | JAETYT010000088.1 | ND |
| Escherichia coli strain ATC-9-6 contig00064, whole genome shotgun sequence | JAETYO010000064.1 | ND |
| Escherichia coli strain ATC-44-40 contig00081, whole genome shotgun sequence | JAETYM010000081.1 | ND |
| Escherichia coli strain ATB47-1 contig00030, whole genome shotgun sequence | JAETYG010000030.1 | lambdoid |
| Escherichia coli strain ATB-15-29 contig00069, whole genome shotgun sequence | JAETYB010000069.1 | ND |
| Escherichia coli strain LSB-2-42b contig00018, whole genome shotgun sequence | JAETYW010000018.1 | Eru8 |
| Escherichia coli strain LSB-2-27b contig00019, whole genome shotgun sequence | JAETYV010000019.1 | Eru8 |
| Escherichia coli strain ATC-B-20-47 contig00012, whole genome shotgun sequence | JAETYP010000012.1 | Eru1 |
| Escherichia coli strain ATC-21-17 contig00069, whole genome shotgun sequence | JAETYJ010000069.1 | ND |
| Escherichia coli strain ATC-11-10 contig00077, whole genome shotgun sequence | JAETYH010000077.1 | ND |
| Escherichia coli strain ATB-21-84 contig00067, whole genome shotgun sequence | JAETYC010000067.1 | ND |
| Escherichia coli strain ATC39-3 contig00011, whole genome shotgun sequence | JAETYR010000011.1 | ND |
| Escherichia coli strain ATB-39-11 contig00011, whole genome shotgun sequence | JAETYE010000011.1 | ND |
| Escherichia coli strain ATB-11-12 contig00046, whole genome shotgun sequence | JAETXZ010000046.1 | Eru10 |

| | | |
|---|---|---|
| Escherichia coli strain LSC6-3 contig00029, whole genome shotgun sequence | JAETZC010000029.1 | Eru10 |
| Escherichia coli strain ATB-23-31c contig00161, whole genome shotgun sequence | JAETYD010000161.1 | ND |
| Escherichia coli strain ATB-10-31 contig00098, whole genome shotgun sequence | JAETXY010000098.1 | ND |
| Escherichia coli strain LSC1-58 contig00138, whole genome shotgun sequence | JAETYZ010000138.1 | ND |
| Escherichia coli strain ATC7-7 contig00075, whole genome shotgun sequence | JAETYU010000075.1 | ND |
| Escherichia coli strain ATC-41-3 contig00056, whole genome shotgun sequence | JAETYL010000056.1 | Eru6 |
| Escherichia coli strain ATC-49-13 contig00211, whole genome shotgun sequence | JAETYN010000211.1 | ND |
| Escherichia coli strain ATB29-1 contig00211, whole genome shotgun sequence | JAETYF010000211.1 | ND |
| Escherichia coli strain LSC1-P21-24 contig00207, whole genome shotgun sequence | JAETZB010000207.1 | ND |
| Escherichia coli strain LSB-P21-51 contig00208, whole genome shotgun sequence | JAETYX010000208.1 | ND |
| Escherichia coli strain LSC1-P21-24 contig00170, whole genome shotgun sequence | JAETZB010000170.1 | ND |
| Escherichia coli strain LSC1-P21-24 contig00095, whole genome shotgun sequence | JAETZB010000095.1 | ND |
| Escherichia coli strain LSB-P21-51 contig00187, whole genome shotgun sequence | JAETYX010000187.1 | ND |
| Escherichia coli strain ATB29-1 contig00181, whole genome shotgun sequence | JAETYF010000181.1 | ND |
| Escherichia coli strain LSB-P21-51 contig00243, whole genome shotgun sequence | JAETYX010000243.1 | ND |
| Escherichia coli strain ATB29-1 contig00102, whole genome shotgun sequence | JAETYF010000102.1 | ND |
| Escherichia coli strain ATC-49-13 contig00089, whole genome shotgun sequence | JAETYN010000089.1 | ND |
| **Stx1** | | |
| Escherichia coli strain ATB6-118 contig00001, whole genome shotgun sequence | JAEUYO010000001.1 | Eru4 |
| Escherichia coli strain LSC1-P21-24 contig00106, whole genome shotgun sequence | JAETZB010000106.1 | ND |
| Escherichia coli strain LSB-P21-51 contig00108, whole genome shotgun sequence | JAETYX010000108.1 | ND |
| Escherichia coli strain ATC-4-67 contig00017, whole genome shotgun sequence | JAETYK010000017.1 | Eru1 |
| Escherichia coli strain ATC-15-17 contig00026, whole genome shotgun sequence | JAETYI010000026.1 | Eru1 |
| Escherichia coli strain ATB29-1 contig00088, whole genome shotgun sequence | JAETYF010000088.1 | ND |
| Escherichia coli strain ATB-23-31c contig00074, whole genome shotgun sequence | JAETYD010000074.1 | ND |
| Escherichia coli strain ATC45-11 contig00091, whole genome shotgun sequence | JAETYS010000091.1 | ND |
| Escherichia coli strain LSC1-7 contig00009, whole genome shotgun sequence | JAETZA010000009.1 | Eru4 |
| Escherichia coli strain LSC1-58 contig00032, whole genome shotgun sequence | JAETYZ010000032.1 | Eru4 |
| Escherichia coli strain LSC-5-20 contig00002, whole genome shotgun sequence | JAETYY010000002.1 | Eru4 |
| Escherichia coli strain ATC7-7 contig00018, whole genome shotgun sequence | JAETYU010000018.1 | Eru4 |
| Escherichia coli strain ATC36-6 contig00045, whole genome shotgun sequence | JAETYQ010000045.1 | Eru4 |
| Escherichia coli strain ATB-14-66 contig00007, whole genome shotgun sequence | JAETYA010000007.1 | Eru4 |
| Escherichia coli strain ATB-10-31 contig00022, whole genome shotgun sequence | JAETXY010000022.1 | Eru4 |
| Escherichia coli strain ATB-23-31c contig00152, whole genome shotgun sequence | JAETYD010000152.1 | ND |
| Escherichia coli strain ATB-23-31c contig00037, whole genome shotgun sequence | JAETYD010000037.1 | |

#### PRJNA680568
#### *Escherichia coli* O80:H2 strains from Switzerland

| Strain | NCBI Accession no | Eru type |
|---|---|---|
| **Stx2** | | |
| Escherichia coli strain 1970-08 1970-08_contig00121, whole genome shotgun sequence | JAEANL010000121.1 | ND |
| Escherichia coli strain S18-215 S18-215_contig00116, whole genome shotgun sequence | JAEANI010000116.1 | ND |

| | | |
|---|---|---|
| Escherichia coli strain S19-18-1 S19-18_contig00081, whole genome shotgun sequence | JAEANE010000081.1 | Eru7 |
| Escherichia coli strain S19-2-2 S19-2_contig00118, whole genome shotgun sequence | JAEAND010000118.1 | ND |
| Escherichia coli strain S19-64-1 S19-64_contig00113, whole genome shotgun sequence | JAEANA010000113.1 | ND |
| Escherichia coli strain S19-677-1 S19-677_contig00129, whole genome shotgun sequence | JAEAMZ010000129.1 | ND |
| Escherichia coli strain 2018-226 STEC2018-226_contig00117, whole genome shotgun sequence | JAEAMV010000117.1 | ND |
| Escherichia coli strain 1384-03 1384-03_contig00133, whole genome shotgun sequence | JAEAMT010000133.1 | ND |
| Escherichia coli strain P17-291 P17-291_contig00069, whole genome shotgun sequence | JAEANK010000069.1 | Eru7 |
| Escherichia coli strain S18-168 S18-168_contig00068, whole genome shotgun sequence | JAEANJ010000068.1 | Eru7 |
| Escherichia coli strain S18-73 S18-73_contig00067, whole genome shotgun sequence | JAEANH010000067.1 | Eru7 |
| Escherichia coli strain S18-9-1 S18-9-1_contig00064, whole genome shotgun sequence | JAEANG010000064.1 | Eru7 |
| Escherichia coli strain S19-101-1 S19-101_contig00070, whole genome shotgun sequence | JAEANF010000070.1 | Eru7 |
| Escherichia coli strain S19-30-1 S19-30_contig00071, whole genome shotgun sequence | JAEANC010000071.1 | Eru7 |
| Escherichia coli strain S19-615-1 S19-615_contig00066, whole genome shotgun sequence | JAEANB010000066.1 | Eru7 |
| Escherichia coli strain S19-710-1 S19-710_contig00078, whole genome shotgun sequence | JAEAMY010000078.1 | Eru7 |
| Escherichia coli strain 2017-299-1 STEC2017-299-1_contig00071, whole genome shotgun sequence | JAEAMX010000071.1 | Eru7 |
| Escherichia coli strain 2017-353-1 STEC2017-353-1_contig00071, whole genome shotgun sequence | JAEAMW010000071.1 | Eru7 |
| Escherichia coli strain 2018-439 STEC2018-439_contig00071, whole genome shotgun sequence | JAEAMU010000071.1 | Eru7 |
| **PRJNA666781**<br>**STEC strains isolated from semi-hard raw milk cheese from Italy** | | |
| **Strain** | **NCBI Accession no** | **Eru type** |
| **Stx2** | | |
| Escherichia coli strain UC4128 STEC_UC4128_contig_75, whole genome shotgun sequence | JACZIB010000075.1 | ND |
| Escherichia coli strain UC4130 STEC_UC4130_contig_74, whole genome shotgun sequence | JACZHZ010000074.1 | ND |
| Escherichia coli strain UC4131 STEC_UC4131_contig_186, whole genome shotgun sequence | JACZHY010000186.1 | ND |
| Escherichia coli strain UC4132 STEC_UC4132_contig_74, whole genome shotgun sequence | JACZHX010000074.1 | ND |
| Escherichia coli strain UC4129 STEC_UC4129_contig_26, whole genome shotgun sequence | JACZIA010000026.1 | ND |
| Escherichia coli strain UC4133 STEC_UC4133_contig_57, whole genome shotgun sequence | JACZHW010000057.1 | ND |
| Escherichia coli strain UC4134 STEC_UC4134_contig_28, whole genome shotgun sequence | JACZHV010000028.1 | ND |
| **Stx1** | | |
| Escherichia coli strain UC4128 STEC_UC4128_contig_17, whole genome shotgun sequence | JACZIB010000017.1 | Eru7 |
| Escherichia coli strain UC4130 STEC_UC4130_contig_17, whole genome shotgun sequence | JACZHZ010000017.1 | Eru7 |
| Escherichia coli strain UC4131 STEC_UC4131_contig_23, whole genome shotgun sequence | JACZHY010000023.1 | Eru7 |
| Escherichia coli strain UC4132 STEC_UC4132_contig_17, whole genome shotgun sequence | JACZHX010000017.1 | Eru7 |
| **PRJNA248042**<br>**National Surveillance of STEC O157:H7 in England** | | |
| **Strain** | **NCBI Accession no** | **Eru type** |
| **Stx2** | | |
| Escherichia coli strain H121560360 SAMN03492009-rid9085603.guided.356, whole genome shotgun sequence | AAVVNG010000004.1 | Eru1 |
| Escherichia coli strain H121560360 SAMN03492009-rid9085603.denovo.333, whole genome shotgun sequence | AAVVVC010000003.1 | Eru1 |

| | | |
|---|---|---|
| Escherichia coli strain H132920427 SAMN03485820-rid9085393.guided.134, whole genome shotgun sequence | AAVVMM010000010.1 | Eru2 |
| Escherichia coli strain H121780072 SAMN03492023-rid9085623.guided.351, whole genome shotgun sequence | AAVVLZ010000015.1 | Eru7 |
| Escherichia coli strain H053000190 SAMN03491912-rid9085593.guided.151, whole genome shotgun sequence | AAVVMI010000016.1 | Eru2 |
| Escherichia coli strain H054440305 SAMN03492026-rid9085653.guided.207, whole genome shotgun sequence | AAVVQR010000015.1 | Eru2 |
| Escherichia coli strain H063100370 SAMN03492028-rid9085673.guided.184, whole genome shotgun sequence | AAVVUX010000016.1 | Eru7 |
| Escherichia coli strain H063100370 SAMN03492028-rid9085673.denovo.134, whole genome shotgun sequence | AAVVMS010000018.1 | Eru7 |
| Escherichia coli strain H094000293 SAMN03492031-rid9085703.guided.352, whole genome shotgun sequence | AAVVOA010000015.1 | Eru7 |
| Escherichia coli strain H102620398 SAMN03492030-rid9085693.guided.275, whole genome shotgun sequence | AAVVVF010000015.1 | Eru7 |
| Escherichia coli strain H094000312 SAMN03492036-rid9085733.guided.276, whole genome shotgun sequence | AAVVXZ010000026.1 | Eru7 |
| Escherichia coli strain H094000312 SAMN03492036-rid9085733.denovo.205, whole genome shotgun sequence | AAVVNS010000026.1 | Eru7 |
| Escherichia coli strain H063160363 SAMN03492043-rid9085743.guided.172, whole genome shotgun sequence | AAVVJN010000025.1 | Eru7 |
| Escherichia coli strain H123780462 SAMN03492044-rid9085793.guided.166, whole genome shotgun sequence | AAVVRM010000028.1 | Eru7 |
| Escherichia coli strain H122440404 SAMN03492048-rid9085833.guided.183, whole genome shotgun sequence | AAVVKN010000025.1 | Eru7 |
| Escherichia coli strain H101980207 SAMN03492054-rid9085873.guided.281, whole genome shotgun sequence | AAVVKJ010000028.1 | Eru7 |
| Escherichia coli strain H043240557 SAMN03492051-rid9085843.guided.166, whole genome shotgun sequence | AAVVVB010000025.1 | Eru7 |
| Escherichia coli strain H123760762 SAMN03492056-rid9085893.guided.229, whole genome shotgun sequence | AAVVTZ010000010.1 | Eru7 |
| Escherichia coli strain H123900440 SAMN03492059-rid9085923.guided.177, whole genome shotgun sequence | AAVVXJ010000022.1 | Eru7 |
| Escherichia coli strain H052720465 SAMN03492060-rid9085933.guided.155, whole genome shotgun sequence | AAVVSU010000026.1 | Eru7 |
| Escherichia coli strain H091920551 SAMN03492063-rid9085963.guided.147, whole genome shotgun sequence | AAVVKR010000027.1 | Eru7 |
| Escherichia coli strain H123800424 SAMN03492066-rid9085993.guided.242, whole genome shotgun sequence | AATHXA010000027.1 | Eru7 |
| Escherichia coli strain H123180903 SAMN03492072-rid9086023.guided.415, whole genome shotgun sequence | AAVVGX010000027.1 | Eru7 |
| Escherichia coli strain H123820316 SAMN03492078-rid9086073.guided.372, whole genome shotgun sequence | AAVVVU010000025.1 | Eru7 |
| Escherichia coli strain H112340317 SAMN03492067-rid9086003.guided.165, whole genome shotgun sequence | AAVVSL010000029.1 | Eru1 |
| Escherichia coli strain H121620527 SAMN03492083-rid9086123.guided.165, whole genome shotgun sequence | AAVVMJ010000033.1 | Eru1 |
| Escherichia coli strain H122880422 SAMN03492081-rid9086103.guided.165, whole genome shotgun sequence | AAVVDW010000029.1 | Eru1 |
| Escherichia coli strain H122460748 SAMN03492087-rid9086143.guided.166, whole genome shotgun sequence | AAVVRE010000037.1 | Eru1 |
| Escherichia coli strain H120540363 SAMN03492095-rid9086193.guided.169, whole genome shotgun sequence | AAVVTU010000036.1 | Eru1 |
| Escherichia coli strain H123400303 SAMN03492107-rid9086273.guided.290, whole genome shotgun sequence | AAVVFF010000037.1 | Eru1 |
| Escherichia coli strain H122920156 SAMN03492121-rid9086373.guided.438, whole genome shotgun sequence | AATHXD010000066.1 | Eru1 |
| Escherichia coli strain H102820427 SAMN03492123-rid9086393.guided.210, whole genome shotgun sequence | AATHXU010000060.1 | Eru1 |
| Escherichia coli strain H121620525 SAMN03492115-rid9086343.guided.168, whole genome shotgun sequence | AAVVTX010000050.1 | Eru5 |
| Escherichia coli strain H093940423 SAMN03492129-rid9086443.guided.195, whole genome shotgun sequence | AAVVMY010000048.1 | Eru7 |
| Escherichia coli strain H132040221 SAMN03492131-rid9086463.guided.169, whole genome shotgun sequence | AAVVTC010000053.1 | Eru1 |
| Escherichia coli strain H123960525 SAMN03492134-rid9086493.guided.198, whole genome shotgun sequence | AAVVLU010000073.1 | Eru1 |

| | | |
|---|---|---|
| Escherichia coli strain H121320380 SAMN03492135-rid9086503.guided.140, whole genome shotgun sequence | AAVVOD010000067.1 | Eru1 |
| Escherichia coli strain H121320380 SAMN03492135-rid9086503.denovo.082, whole genome shotgun sequence | AATHWY010000053.1 | Eru7 |
| Escherichia coli strain H122900266 SAMN03492148-rid9086603.guided.215, whole genome shotgun sequence | AAVVQZ010000056.1 | Eru7 |
| Escherichia coli strain H123480455 SAMN03492142-rid9086553.guided.197, whole genome shotgun sequence | AAVVED010000052.1 | Eru5 |
| Escherichia coli strain H123740302 SAMN03492155-rid9086673.guided.274, whole genome shotgun sequence | AAVVTC010000107.1 | Eru1 |
| Escherichia coli strain H123540543 SAMN03492158-rid9086703.guided.208, whole genome shotgun sequence | AAVVWC010000072.1 | Eru7 |
| Escherichia coli strain H121600272 SAMN03492156-rid9086683.guided.223, whole genome shotgun sequence | AAVVIM010000068.1 | Eru7 |
| Escherichia coli strain H123680777 SAMN03492169-rid9086783.guided.153, whole genome shotgun sequence | AAVVJS010000064.1 | Eru1 |
| Escherichia coli strain H121980151 SAMN03492170-rid9086793.guided.159, whole genome shotgun sequence | AAVVFK010000056.1 | Eru1 |
| Escherichia coli strain H122860601 SAMN03492171-rid9086803.guided.179, whole genome shotgun sequence | AAVVXH010000123.1 | Eru7 |
| Escherichia coli strain H122920157 SAMN03492172-rid9086813.guided.176, whole genome shotgun sequence | AAVVEV010000056.1 | Eru1 |
| Escherichia coli strain H134240606 SAMN03492175-rid9086843.guided.135, whole genome shotgun sequence | AAVVCY010000055.1 | Eru1 |
| Escherichia coli strain H134240606 SAMN03492175-rid9086843.denovo.113, whole genome shotgun sequence | AAVVFQ010000059.1 | Eru1 |
| Escherichia coli strain H122220862 SAMN03492174-rid9086833.guided.158, whole genome shotgun sequence | AAVVPS010000058.1 | Eru1 |
| Escherichia coli strain H131980146 SAMN03492180-rid9086873.guided.160, whole genome shotgun sequence | AAVVJC010000072.1 | Eru1 |
| Escherichia coli strain H122140365 SAMN03492185-rid9086923.guided.176, whole genome shotgun sequence | AAVVKG010000054.1 | Eru2 |
| Escherichia coli strain H121100178 SAMN03492188-rid9086953.guided.106, whole genome shotgun sequence | AAVVPA010000065.1 | Eru1 |
| Escherichia coli strain H121100178 SAMN03492188-rid9086953.denovo.051, whole genome shotgun sequence | AAVVHE010000064.1 | Eru1 |
| Escherichia coli strain H121620528 SAMN03492190-rid9086973.guided.173, whole genome shotgun sequence | AAVVHY010000075.1 | ND |
| Escherichia coli strain H093700590 SAMN03492192-rid9086993.guided.171, whole genome shotgun sequence | AAVVGZ010000088.1 | ND |
| Escherichia coli strain H122460420 SAMN03492195-rid9087023.guided.248, whole genome shotgun sequence | AAVVRJ010000077.1 | Eru7 |
| Escherichia coli strain H121820144 SAMN03492208-rid9087123.guided.155, whole genome shotgun sequence | AAVVWR010000124.1 | Eru2 |
| **Stx1** | | |
| Escherichia coli strain H125180252 SAMN03492074-rid9086033.denovo.004, whole genome shotgun sequence | AAVVXL010000004.1 | eru4 |
| Escherichia coli strain E1653600 SAMN03703076-rid9072093.denovo.004, whole genome shotgun sequence | AATHWC010000004.1 | Eru13 |
| Escherichia coli strain H131620832 SAMN03492568-rid9038543.denovo.007, whole genome shotgun sequence | AAVVNQ010000007.1 | lambdoid |
| Escherichia coli strain H132360377 SAMN03492323-rid9087503.denovo.001, whole genome shotgun sequence | AAVVSB010000001.1 | lambdoid |
| Escherichia coli strain H133040516 SAMN03492437-rid9037593.denovo.004, whole genome shotgun sequence | AAVVPQ010000004.1 | eru4 |
| Escherichia coli strain H103460518 SAMN03492465-rid9037983.denovo.006, whole genome shotgun sequence | AAVVOT010000006.1 | lambdoid |
| Escherichia coli strain H113320446 SAMN03492141-rid9086543.denovo.064, whole genome shotgun sequence | AAVVVG010000064.1 | Eru1 |
| Escherichia coli strain H130960185 SAMN03492409-rid9087913.denovo.075, whole genome shotgun sequence | AAVVQM010000074.1 | Eru1 |
| Escherichia coli strain E113096 SAMN03703088-rid9072583.denovo.051, whole genome shotgun sequence | AATHVY010000051.1 | lambdoid |
| Escherichia coli strain E1688150 SAMN03702926-rid9033583.denovo.059, whole genome shotgun sequence | AATIBF010000058.1 | lambdoid |
| Escherichia coli strain H103780083 SAMN03492786-rid9061363.denovo.051, whole genome shotgun sequence | AAVVJM010000050.1 | Eru11 |

| | | |
|---|---|---|
| Escherichia coli strain H122220222 SAMN03492104-rid9086243.denovo.054, whole genome shotgun sequence | AAVVWI010000053.1 | lambdoid |
| Escherichia coli strain H123000125 SAMN03492075-rid9086043.denovo.061, whole genome shotgun sequence | AAVVXF010000060.1 | lambdoid |
| Escherichia coli strain H134660555 SAMN03702957-rid9033863.denovo.051, whole genome shotgun sequence | AATIAB010000050.1 | lambdoid |
| Escherichia coli strain H122980845 SAMN03492153-rid9086653.denovo.106, whole genome shotgun sequence | AAVVUY010000106.1 | lambdoid |
| Escherichia coli strain WX011665S01E SAMN03496101-rid9061543.denovo.055, whole genome shotgun sequence | AAVVGH010000054.1 | lambdoid |
| Escherichia coli strain H123340502 SAMN03492674-rid9039133.denovo.051, whole genome shotgun sequence | AAVVMC010000050.1 | lambdoid |
| Escherichia coli strain H122960360 SAMN03492359-rid9087693.denovo.092, whole genome shotgun sequence | AAVVRI010000092.1 | lambdoid |
| Escherichia coli strain H123980865 SAMN03492202-rid9087063.denovo.054, whole genome shotgun sequence | AAVVTR010000053.1 | lambdoid |
| Escherichia coli strain H122760538 SAMN03492653-rid9038943.denovo.066, whole genome shotgun sequence | AAVVMW010000065.1 | lambdoid |
| Escherichia coli strain WX016375S01E SAMN03496188-rid9080733.denovo.051, whole genome shotgun sequence | AAVVDM010000050.1 | lambdoid |
| Escherichia coli strain WX017993S01E SAMN03496184-rid9080543.denovo.052, whole genome shotgun sequence | AAVVEW010000051.1 | lambdoid |
| Escherichia coli strain H123040570 SAMN03492632-rid9038833.denovo.093, whole genome shotgun sequence | AAVVHC010000093.1 | lambdoid |
| Escherichia coli strain H123120512 SAMN03492068-rid9086013.denovo.074, whole genome shotgun sequence | AAVVXJ010000073.1 | lambdoid |
| Escherichia coli strain H123200240 SAMN03492225-rid9087283.denovo.052, whole genome shotgun sequence | AAVVSS010000051.1 | lambdoid |
| Escherichia coli strain H124440480 SAMN03492126-rid9086423.denovo.052, whole genome shotgun sequence | AAVVVT010000051.1 | ND |
| Escherichia coli strain H104620471 SAMN03492154-rid9086663.denovo.062, whole genome shotgun sequence | AAVVUS010000061.1 | lambdoid |
| Escherichia coli strain H102980650 SAMN03492463-rid9037963.denovo.060, whole genome shotgun sequence | AAVVOS010000060.1 | lambdoid |
| Escherichia coli strain H132080461 SAMN03492034-rid9085713.denovo.065, whole genome shotgun sequence | AAVVYL010000064.1 | lambdoid |
| Escherichia coli strain H122440402 SAMN03492475-rid9038083.denovo.055, whole genome shotgun sequence | AAVVOH010000054.1 | lambdoid |
| Escherichia coli strain H121800664 SAMN03702960-rid9033893.denovo.060, whole genome shotgun sequence | AATIAA010000059.1 | lambdoid |
| Escherichia coli strain H113580158 SAMN03492105-rid9086253.denovo.052, whole genome shotgun sequence | AAVVWF010000051.1 | lambdoid |
| Escherichia coli strain H121380674 SAMN03492111-rid9086303.denovo.068, whole genome shotgun sequence | AAVVWD010000068.1 | lambdoid |
| Escherichia coli strain WX016320S01E SAMN03496190-rid9080753.denovo.050, whole genome shotgun sequence | AAVVDJ010000049.1 | lambdoid |
| Escherichia coli strain H123740300 SAMN03492626-rid9038773.denovo.058, whole genome shotgun sequence | AAVVHJ010000057.1 | lambdoid |
| Escherichia coli strain E1753820 SAMN03703008-rid9034313.denovo.054, whole genome shotgun sequence | AATHYI010000053.1 | lambdoid |
| Escherichia coli strain H121620536 SAMN03492418-rid9088013.denovo.089, whole genome shotgun sequence | AAVVQC010000089.1 | lambdoid |
| Escherichia coli strain H130740146 SAMN03492441-rid9037623.denovo.054, whole genome shotgun sequence | AAVVPM010000053.1 | lambdoid |
| Escherichia coli strain H132480192 SAMN03492045-rid9085803.denovo.070, whole genome shotgun sequence | AAVVYG010000069.1 | lambdoid |
| Escherichia coli strain H123880634 SAMN03492052-rid9085853.denovo.096, whole genome shotgun sequence | AAVVYD010000096.1 | lambdoid |
| Escherichia coli strain H122980190 SAMN03492442-rid9037633.denovo.124, whole genome shotgun sequence | AAVVPO010000124.1 | lambdoid |
| Escherichia coli strain H123840423 SAMN03492519-rid9038683.denovo.069, whole genome shotgun sequence | AAVVHS010000069.1 | lambdoid |
| Escherichia coli strain H121100178 SAMN03492188-rid9086953.denovo.054, whole genome shotgun sequence | AAVVTX010000053.1 | lambdoid |
| Escherichia coli strain H132940671 SAMN03492649-rid9038913.denovo.083, whole genome shotgun sequence | AAVVNC010000081.1 | Eru1 |
| Escherichia coli strain H133040516 SAMN03492437-rid9037593.guided.073, whole genome shotgun sequence | AAVVPQ010000128.1 | Eru4 |

| | | |
|---|---|---|
| Escherichia coli strain E1745770 SAMN03702990-rid9034143.denovo.061, whole genome shotgun sequence | AATHZE010000060.1 | ND |
| Escherichia coli strain E1747730 SAMN03702922-rid9033553.denovo.063, whole genome shotgun sequence | AATIBI010000062.1 | lambdoid |
| Escherichia coli strain H130300235 SAMN03492425-rid9033073.denovo.183, whole genome shotgun sequence | AAVVQB010000182.1 | ND |
| **PRJNA715185<br>Diversity of STEC in flour from Germany** | | |
| **Strain** | **NCBI Accession no** | **Eru type** |
| **Stx2** | | |
| Escherichia coli strain BfR-EC-17777 60, whole genome shotgun sequence | JAGEWD010000060.1 | ND |
| Escherichia coli strain BfR-EC-17374 67, whole genome shotgun sequence | JAGEXU010000067.1 | ND |
| Escherichia coli strain BfR-EC-17741 53, whole genome shotgun sequence | JAGEWM010000053.1 | ND |
| Escherichia coli strain BfR-EC-17656 61, whole genome shotgun sequence | JAGEXI010000061.1 | ND |
| Escherichia coli strain BfR-EC-17705 59, whole genome shotgun sequence | JAGEWZ010000059.1 | ND |
| Escherichia coli strain BfR-EC-17730 55, whole genome shotgun sequence | JAGEWO010000055.1 | ND |
| Escherichia coli strain BfR-EC-17709 60, whole genome shotgun sequence | JAGEWY010000060.1 | ND |
| Escherichia coli strain BfR-EC-17710 59, whole genome shotgun sequence | JAGEWX010000059.1 | ND |
| Escherichia coli strain BfR-EC-17722 59, whole genome shotgun sequence | JAGEWS010000059.1 | ND |
| Escherichia coli strain BfR-EC-17655 58, whole genome shotgun sequence | JAGEXJ010000058.1 | ND |
| Escherichia coli strain BfR-EC-17679 44, whole genome shotgun sequence | JAGEXB010000044.1 | Eru9 |
| Escherichia coli strain BfR-EC-17841 contig00037, whole genome shotgun sequence | JAGEVW010000037.1 | Eru9 |
| Escherichia coli strain BfR-EC-17718 47, whole genome shotgun sequence | JAGEWW010000047.1 | ND |
| Escherichia coli strain BfR-EC-17719 47, whole genome shotgun sequence | JAGEWV010000047.1 | ND |
| Escherichia coli strain BfR-EC-17680 51, whole genome shotgun sequence | JAGEXA010000051.1 | Eru9 |
| Escherichia coli strain BfR-EC-17678 39, whole genome shotgun sequence | JAGEXC010000039.1 | Eru9 |
| Escherichia coli strain BfR-EC-17778 35, whole genome shotgun sequence | JAGEWC010000035.1 | ND |
| Escherichia coli strain BfR-EC-17779 34, whole genome shotgun sequence | JAGEWB010000034.1 | ND |
| Escherichia coli strain BfR-EC-17387 9, whole genome shotgun sequence | JAGEXQ010000009.1 | Eru9 |
| **Stx1** | | |
| Escherichia coli strain BfR-EC-17379 contig00014, whole genome shotgun sequence | JAGEXT010000014.1 | ND |
| Escherichia coli strain BfR-EC-17386 contig00011, whole genome shotgun sequence | JAGEXR010000011.1 | ND |
| Escherichia coli strain BfR-EC-17751 contig00012, whole genome shotgun sequence | JAGEWI010000012.1 | ND |
| Escherichia coli strain BfR-EC-17856 contig00002, whole genome shotgun sequence | JAGEVS010000002.1 | ND |
| Escherichia coli strain BfR-EC-17721 contig00001, whole genome shotgun sequence | JAGEWT010000001.1 | ND |
| Escherichia coli strain BfR-EC-17380 contig00001, whole genome shotgun sequence | JAGEXS010000001.1 | ND |
| **PRJNA643688<br>Genome Sequences of twelve Shiga Toxin-Producing *Escherichia coli* Strains Isolated from dairy cattle in Portugal** | | |
| **Strain** | **NCBI Accession no** | **Eru type** |
| **Stx2** | | |
| Escherichia coli strain E7N3P7C8A Contig_26_consensus_sequence, whole genome shotgun sequence | JACBWM010000026.1 | Eru1 |

| | | |
|---|---|---|
| Escherichia coli strain E7N8P4C1 Contig_42_consensus_sequence, whole genome shotgun sequence | JACBWI010000042.1 | ND |
| Escherichia coli strain E7N15P4C10 Contig_2_consensus_sequence, whole genome shotgun sequence | JACBWG010000002.1 | Eru7 |
| Escherichia coli strain E7N18P5C8G Contig_17_consensus_sequence, whole genome shotgun sequence | JACBWE010000017.1 | Eru8 |
| Escherichia coli strain E7N6P4C8A Contig_40_consensus_sequence, whole genome shotgun sequence | JACBWL010000040.1 | ND |
| Escherichia coli strain E7N6P4C8C Contig_53_consensus_sequence, whole genome shotgun sequence | JACBWK010000053.1 | ND |
| Escherichia coli strain E7N6P4C8F Contig_38_consensus_sequence, whole genome shotgun sequence | JACBWJ010000038.1 | ND |
| Escherichia coli strain E7N16P4C9 Contig_58_consensus_sequence, whole genome shotgun sequence | JACBWF010000058.1 | ND |
| Escherichia coli strain E7N6P4C8A Contig_34_consensus_sequence, whole genome shotgun sequence | JACBWL010000034.1 | ND |
| Escherichia coli strain E7N16P4C9 Contig_43_consensus_sequence, whole genome shotgun sequence | JACBWF010000043.1 | ND |
| Escherichia coli strain E7N8P4C1 Contig_5_consensus_sequence, whole genome shotgun sequence | JACBWI010000005.1 | Eru8 |
| Escherichia coli strain E7N6P4C8C Contig_36_consensus_sequence, whole genome shotgun sequence | JACBWK010000036.1 | ND |
| Escherichia coli strain E7N6P4C8C Contig_59_consensus_sequence, whole genome shotgun sequence | JACBWK010000059.1 | ND |
| Escherichia coli strain E7N16P4C9 Contig_59_consensus_sequence, whole genome shotgun sequence | JACBWF010000059.1 | ND |
| Escherichia coli strain E7N6P4C8F Contig_63_consensus_sequence, whole genome shotgun sequence | JACBWJ010000063.1 | ND |
| **Stx1** | | |
| Escherichia coli strain E7N18P5C8G Contig_11_consensus_sequence, whole genome shotgun sequence | JACBWE010000011.1 | Eru7 |
| Escherichia coli strain E7N6P4C8A Contig_12_consensus_sequence, whole genome shotgun sequence | JACBWL010000012.1 | Eru6 |
| Escherichia coli strain E7N6P4C8C Contig_24_consensus_sequence, whole genome shotgun sequence | JACBWK010000024.1 | Eru6 |
| Escherichia coli strain E7N6P4C8F Contig_36_consensus_sequence, whole genome shotgun sequence | JACBWJ010000036.1 | ND |
| Escherichia coli strain E7N16P4C9 Contig_28_consensus_sequence, whole genome shotgun sequence | JACBWF010000028.1 | Eru6 |
| Escherichia coli strain E7V12P1C4G Contig_79_consensus_sequence, whole genome shotgun sequence | JACBWN010000079.1 | ND |
| Escherichia coli strain E7N3P7C8A Contig_56_consensus_sequence, whole genome shotgun sequence | JACBWM010000056.1 | lambdoid |
| Escherichia coli strain E7V3P1C1 Contig_7_consensus_sequence, whole genome shotgun sequence | JACBWP010000007.1 | ND |
| Escherichia coli strain E7V4P1C10 Contig_30_consensus_sequence, whole genome shotgun sequence | JACBWO010000030.1 | ND |
| Escherichia coli strain E7N12P2C4 Contig_71_consensus_sequence, whole genome shotgun sequence | JACBWH010000071.1 | ND |
| **PRJNA438214**<br>**Draft genomes sequences of 18 *Escherichia coli* STEC strains from Switzerland** | | |
| **Strain** | **NCBI Accession no** | **Eru type** |
| **Stx2** | | |
| Escherichia coli strain 364060-17 NODE_89_length_17522_cov_9.483185, whole genome shotgun sequence | PYSE01000089.1 | Eru7 |
| Escherichia coli strain 364062-17 NODE_5_length_99620_cov_14.092529, whole genome shotgun sequence | PYSC01000005.1 | Eru7 |
| Escherichia coli strain 364064-17 NODE_95_length_16865_cov_12.828414, whole genome shotgun sequence | PYSA01000095.1 | Eru7 |
| Escherichia coli strain 364068-17 NODE_93_length_15220_cov_19.738422, whole genome shotgun sequence | PYRX01000093.1 | Eru7 |
| Escherichia coli strain 364073-17 NODE_64_length_24383_cov_39.192117, whole genome shotgun sequence | PYRU01000064.1 | Eru7 |
| Escherichia coli strain 364082-17 NODE_100_length_13439_cov_15.958609, whole genome shotgun sequence | PYRR01000100.1 | Eru7 |
| Escherichia coli strain 364072-17 scaffold_0186, whole genome shotgun sequence | PYRP01000186.1 | ND |
| Escherichia coli strain 364079-17 scaffold_0171, whole genome shotgun sequence | PYRO01000171.1 | ND |

| | | |
|---|---|---|
| Escherichia coli strain 364059-17 NODE_126_length_11966_cov_12.106850, whole genome shotgun sequence | PYSF01000126.1 | Eru7 |
| Escherichia coli strain 364061-17 NODE_88_length_15513_cov_19.573573, whole genome shotgun sequence | PYSD01000088.1 | Eru7 |
| Escherichia coli strain 364063-17 scaffold_0125, whole genome shotgun sequence | PYSB01000125.1 | Eru7 |
| Escherichia coli strain 364067-17 scaffold_0011, whole genome shotgun sequence | PYRY01000011.1 | Eru7 |
| Escherichia coli strain 364069-17 NODE_71_length_15513_cov_65.140517, whole genome shotgun sequence | PYRW01000071.1 | Eru7 |
| Escherichia coli strain 364070-17 scaffold_0017, whole genome shotgun sequence | PYRV01000017.1 | Eru7 |
| Escherichia coli strain 364075-17 NODE_73_length_15513_cov_42.632068, whole genome shotgun sequence | PYRT01000073.1 | Eru7 |
| Escherichia coli strain 364077-17 NODE_88_length_15513_cov_17.408748, whole genome shotgun sequence | PYRS01000088.1 | Eru7 |
| Escherichia coli strain 364071-17 scaffold_0089, whole genome shotgun sequence | PYRQ01000089.1 | Eru7 |
| Escherichia coli strain 364066-17 scaffold_0737, whole genome shotgun sequence | PYRZ01000737.1 | ND |
| Escherichia coli strain 364066-17 scaffold_0504, whole genome shotgun sequence | PYRZ01000504.1 | ND |

Table S3

Information about the 260 CI sequences used in phylogenetic analysis

| CI ID (NCBI) | Phage/Strain | Serotype | NCBI ACCESSION NO | Origin | Year | Stx type | Eru type |
|---|---|---|---|---|---|---|---|
| ACI32363.1 | Enterobacteria phage YYZ-2008 | O157:H7 | FJ184280 | Canada | 2008 | Stx1 | Eru2 |
| ACI43117.1 | Stx2 Converting phage 1717 | O157:H7 | FJ188381 | Canada | 2008 | Stx2 | Eru2 |
| ADN68413.1 | Stx2 converting phage vB_EcoP_24B | O157:H7 | HM208303.1 | UK | 2010 | Stx2 | Eru5 |
| AHZ95179.1 | Shigella phage POCJ13 | | KJ603229.1 | USA | 2014 | Stx1 | Eru4 |
| AIF74338.1 | Escherichia Stx1-converting recombinant phage HUN/2013 | O157:H7 | KJ909655.1 | Hungary | 2013 | Stx1 | Eru3 |
| AKI86003.1 | Escherichia phage PA4 | O157:H7 | KP682372.1 | USA | 2015 | Stx2 | Eru3 |
| AKI86096.1 | Escherichia phage PA5 | O157:H7 | KP682373.1 | USA | 2015 | Stx2 | Eru3 |
| AKI86274.1 | Escherichia phage PA11 | O157:H7 | KP682375.1 | USA | 2015 | Stx2 | Eru3 |
| AKI86367.1 | Escherichia phage PA16 | O157:H7 | KP682377.1 | USA | 2015 | Stx2 | Eru3 |
| AKI86553.1 | Escherichia phage PA21 | O157:H7 | KP682379.1 | USA | 2015 | Stx2 | Eru3 |
| AKI86648.1 | Escherichia phage PA27 | O157:H7 | KP682380.1 | USA | 2015 | Stx2 | Eru3 |
| AKI86748.1 | Escherichia phage PA28 | O157:H7 | KP682381.1 | USA | 2015 | Stx2 | lambdoid |
| AKI86834.1 | Escherichia phage PA29 | O157:H7 | KP682382.1 | USA | 2015 | Stx2 | Eru3 |
| AKI87206.1 | Escherichia phage PA36 | O157:H7 | KP682386.1 | USA | 2015 | Stx2 | Eru3 |
| AKI87302.1 | Escherichia phage PA42 | O157:H7 | KP682387.1 | USA | 2015 | Stx2 | Eru3 |
| AKI87492.1 | Escherichia phage PA45 | O157:H7 | KP682389.1 | USA | 2015 | Stx2 | Eru3 |
| AKI87588.1 | Escherichia phage PA50 | O157:H7 | KP682390.1 | USA | 2015 | Stx2 | Eru3 |
| AKI87680.1 | Escherichia phage PA52 | O157:H7 | KP682392.1 | USA | 2015 | Stx2 | Eru3 |
| AKJ74710.1 | Escherichia phage PA12 | O157:H7 | KP682376.1 | USA | 2015 | Stx2 | Eru3 |
| ANJ63820.1 | Stx1 converting phage AU5Stx1 | O157 | KU977419.1 | Australia | 2016 | Stx1 | lambdoid |
| ANJ63898.1 | Stx1 converting phage AU6Stx1 | O157 | KU977420.1 | Australia | 2016 | Stx1 | lambdoid |
| AVD99089.1 | Escherichia phage GER2 | O117:H7 | MG710528.1 | UK | 2017 | Stx1 | Eru1 |
| BAB87968.1 | Stx2 Converting phage I | O157:H7 | AP004402 | Japan | 2001 | Stx2 | lambdoid |
| BAC77937.1 | Morioka V526 | O157:H7 | AP005153.1 | Japan | 2003 | Stx1 | Eru3 |
| BAC78103.1 | Morioka V526 | O157:H7 | AP005154.1 | Japan | | Stx2 | Eru3 |
| BAT31827.1 | Stx2-converting phage Stx2a_F403 proviral | O157:H7 | AP012529.1 | Japan | 2012 | Stx2 | Eru5 |
| BAT31911.1 | Stx2-converting phage Stx2a_F349 | O157:H7 | AP012530.1 | Japan | 2015 | Stx2 | Eru2 |
| BAT32000.1 | Stx2-converting phage Stx2a_F422 proviral | O157:H7 | AP012531.1 | Japan | 2012 | Stx2 | lambdoid |
| BAT32093.1 | Stx2-converting phage Stx2a_F451 | O157:H8 | AP012532.1 | Japan | 2012 | Stx2 | Eru5 |
| BAT32177.1 | Stx2-converting phage Stx2a_F723 | O157:H9 | AP012533.1 | Japan | 2012 | Stx2 | lambdoid |
| BAT32263.1 | Stx2-converting phage Stx2a_F765 proviral | O157:H7 | AP012534.1 | Japan | 2012 | Stx2 | Eru1 |
| BAT32321.1 | Stx2-converting phage Stx2a_WGPS9 proviral | O157:H7 | AP012535.1 | Japan | 2012 | Stx2 | lambdoid |
| BAT32387.1 | Stx2-converting phage Stx2a_1447 proviral | O157:H7 | AP012536.1 | Japan | 2012 | Stx2 | Eru6 |
| BAT32452.1 | Stx2-converting phage Stx2a_WGPS2 proviral | O157:H7 | AP012537.1 | Japan | 2012 | Stx2 | Eru6 |
| BAT32531.1 | Stx2-converting phage Stx2a_WGPS4 proviral | O157:H7 | AP012538.1 | Japan | 2012 | Stx2 | Eru2 |

| | | | | | | | |
|---|---|---|---|---|---|---|---|
| BAT32614.1 | Stx2-converting phage Stx2a_WGPS6 proviral | O157:H7 | AP012539.1 | Japan | 2012 | Stx2 | Eru2 |
| BAT32693.1 | Stx2-converting phage Stx2a_WGPS8 proviral | O157:H7 | AP012540.1 | Japan | 2012 | Stx2 | Eru2 |
| BCI48989.1 | Stx2a-converting phage Stx2_14040 | O145:H28 | LC567818.1 | Japan | 2020 | Stx2 | Eru7 |
| BCI49050.1 | Stx1a-converting phage Stx1_14040 | O145:H28 | LC567819.1 | Japan | 2020 | Stx1 | Eru7 |
| BCI49153.1 | Stx2a-converting phage Stx2_14744 | O145:H28 | LC567820.1 | Japan | 2020 | Stx2 | Eru7 |
| BCI49214.1 | Stx1a-converting phage Stx1_14744 | O145:H28 | LC567821.1 | Japan | 2020 | Stx1 | Eru7 |
| CAC83529.1 | Enterobacteria phage phiP27 | ONT:H- | AJ298298 | Germany | 2002 | Stx2 | Eru7 |
| CAD88825.1 | Phage BP-4795 complete genome | O84:H4 | AJ556162.1 | Germany | 2003 | Stx1 | lambdoid |
| CAQ82008.1 | Enterobacteria phage 2851 | O157:H7 | FM180578.1 | Germany | 1993 | Stx2 | Eru2 |
| CCG06176.1 | Escherichia phage P13374 proviral | O104:H4 | HE664024.1 | Germany | 2011 | Stx2 | Eru1 |
| CDK12686.1 | Escherichia phage P13803 | O2:H27 | HG792102.1 | Germany | 2013 | Stx2 | Eru1 |
| CDK23993.1 | Escherichia phage P14437 | O104:H4 | HG792105.1 | Norway | 2006 | Stx2 | Eru1 |
| CDL18842.1 | Escherichia phage P13771 | O104:H4 | HG792104.1 | Germany | 2009 | Stx2 | Eru1 |
| EEC26960.1 | TW14588 | O157:H7 | ABKY02000003 | USA | 2006 | Stx2 | Eru3 |
| EEC27821.1 | TW14588 | O157:H7 | ABKY02000002 | USA | 2006 | Stx2 | Eru1 |
| EFL4537683.1 | E1653600 | O157:H7 | AATHWC010000004.1 | UK | | Stx1 | Eru13 |
| EFL4561882.1 | E113096 | O157:H7 | AATHVY010000051.1 | UK | | Stx1 | lambdoid |
| EFL4759493.1 | H123800424 | O157:H7 | AATHXA010000027.1 | UK | | Stx2 | Eru7 |
| EFL4780259.1 | H122920156 | O157:H7 | AATHXD010000066.1 | UK | | Stx2 | Eru1 |
| EFL4785786.1 | H121320380 | O157:H7 | AATHWY010000053.1 | UK | | Stx2 | Eru7 |
| EFL4810920.1 | H102820427 | O157:H7 | AATHXU010000060.1 | UK | | Stx2 | Eru1 |
| EFL4852155.1 | E1753820 | O157:H7 | AATHYI010000053.1 | UK | | Stx1 | lambdoid |
| EFL5060758.1 | H121800664 | O157:H7 | AATIAA010000059.1 | UK | | Stx1 | lambdoid |
| EFL5163126.1 | H134660555 | O157:H7 | AATIAB010000050.1 | UK | | Stx1 | lambdoid |
| EFL5223636.1 | E1688150 | O157:H7 | AATIBF010000058.1 | UK | | Stx1 | lambdoid |
| EGE5688604.1 | H134240606 | O157:H7 | AAVVCY010000055.1 | UK | | Stx2 | Eru2 |
| EGE5822072.1 | WX016320S01E | O157:H7 | AAVVDJ010000049.1 | UK | | Stx1 | lambdoid |
| EGE5827186.1 | WX016375S01E | O157:H7 | AAVVDM010000050.1 | UK | | Stx1 | lambdoid |
| EGE5841739.1 | H122880422 | O157:H7 | AAVVDW010000029.1 | UK | | Stx2 | Eru1 |
| EGE5909574.1 | H123480455 | O157:H7 | AAVVED010000052.1 | UK | | Stx2 | Eru5 |
| EGE5954436.1 | H122920157 | O157:H7 | AAVVEV010000056.1 | UK | | Stx2 | Eru1 |
| EGE5963960.1 | H123400303 | O157:H7 | AAVVFF010000037.1 | UK | | Stx2 | Eru1 |
| EGE6051643.1 | WX017993S01E | O157:H7 | AAVVEW010000051.1 | UK | | Stx1 | lambdoid |
| EGE6056760.1 | H134240606 | O157:H7 | AAVVFQ010000059.1 | UK | | Stx2 | Eru1 |
| EGE6062049.1 | H121980151 | O157:H7 | AAVVFK010000056.1 | UK | | Stx2 | Eru1 |
| EGE6148684.1 | WX011665S01E | O157:H7 | AAVVGH010000054.1 | UK | | Stx1 | lambdoid |
| EGE6228835.1 | H123180903 | O157:H7 | AAVVGX010000027.1 | UK | | Stx2 | Eru7 |
| EGE6276458.1 | H121100178 | O157:H7 | AAVVHE010000064.1 | UK | | Stx2 | Eru1 |
| EGE6286513.1 | H123040570 | O157:H7 | AAVVHC010000093.1 | UK | | Stx1 | lambdoid |
| EGE6307131.1 | H123740300 | O157:H7 | AAVVHJ010000057.1 | UK | | Stx1 | lambdoid |

| | | | | | | | |
|---|---|---|---|---|---|---|---|
| EGE6419498.1 | H123840423 | O157:H7 | AAVVHS010000069.1 | UK | | Stx1 | lambdoid |
| EGE6460037.1 | H121600272 | O157:H7 | AAVVIM010000068.1 | UK | | Stx2 | Eru7 |
| EGE6542355.1 | H131980146 | O157:H7 | AAVVJC010000072.1 | UK | | Stx2 | Eru1 |
| EGE6605736.1 | H063160363 | O157:H7 | AAVVJN010000025.1 | UK | | Stx2 | Eru7 |
| EGE6633054.1 | H123680777 | O157:H7 | AAVVJS010000064.1 | UK | | Stx2 | Eru1 |
| EGE6689883.1 | H122140365 | O157:H7 | AAVVKG010000054.1 | UK | | Stx2 | Eru2 |
| EGE6708400.1 | H101980207 | O157:H7 | AAVVKJ010000028.1 | UK | | Stx2 | Ery7 |
| EGE6722855.1 | H122440404 | O157:H7 | AAVVKN010000025.1 | UK | | Stx2 | Eru7 |
| EGE6749235.1 | H091920551 | O157:H7 | AAVVKR010000027.1 | UK | | Stx2 | Eru7 |
| EGE6922782.1 | H121780072 | O157:H7 | AAVVLZ010000015.1 | UK | | Stx2 | Eru7 |
| EGE6963583.1 | H121620527 | O157:H7 | AAVVMJ010000033.1 | UK | | Stx2 | Eru1 |
| EGE6977926.1 | H053000190 | O157:H7 | AAVVMI010000016.1 | UK | | Stx2 | Eru2 |
| EGE6982261.1 | H132920427 | O157:H7 | AAVVMM010000010.1 | UK | | Stx2 | Eru2 |
| EGE7018974.1 | H063100370 | O157:H7 | AAVVMS010000018.1 | UK | | Stx2 | Eru7 |
| EGE7050911.1 | H093940423 | O157:H7 | AAVVMY010000048.1 | UK | | Stx2 | Eru7 |
| EGE7055905.1 | H122760538 | O157:H7 | AAVVMW010000065.1 | UK | | Stx1 | lambdoid |
| EGE7073250.1 | H121560360 | O157:H7 | AAVVNG010000004.1 | UK | | Stx2 | Eru1 |
| EGE7134837.1 | H131620832 | O157:H7 | AAVVNQ010000007.1 | UK | | Stx1 | lambdoid |
| EGE7146195.1 | H094000312 | O157:H7 | AAVVNS010000026.1 | UK | | Stx2 | Eru7 |
| EGE7166646.1 | H094000293 | O157:H7 | AAVVOA010000015.1 | UK | | Stx2 | Eru7 |
| EGE7199773.1 | H121320380 | O157:H7 | AAVVOD010000067.1 | UK | | Stx2 | Eru1 |
| EGE7230283.1 | H122440402 | O157:H7 | AAVVOH010000054.1 | UK | | Stx1 | lambdoid |
| EGE7333290.1 | H121100178 | O157:H7 | AAVVPA010000065.1 | UK | | Stx2 | Eru1 |
| EGE7343382.1 | H130740146 | O157:H7 | AAVVPM010000053.1 | UK | | Stx1 | lambdoid |
| EGE7370760.1 | H133040516 | O157:H7 | AAVVPQ010000004.1 | UK | | Stx1 | Eru4 |
| EGE7389752.1 | H122220862 | O157:H7 | AAVVPS010000058.1 | UK | | Stx2 | Eru1 |
| EGE7394257.1 | H122980190 | O157:H7 | AAVVPO010000124.1 | UK | | Stx1 | lambdoid |
| EGE7456400.1 | H121620536 | O157:H7 | AAVVQC010000089.1 | UK | | Stx1 | lambdoid |
| EGE7512252.1 | H130960185 | O157:H7 | AAVVQM010000074.1 | UK | | Stx1 | Eru1 |
| EGE7526414.1 | H123880824 | O157:H7 | AAVVQR010000015.1 | UK | | Stx2 | Eru2 |
| EGE7568803.1 | H122900266 | O157:H7 | AAVVQZ010000056.1 | UK | | Stx2 | Eru7 |
| EGE7603615.1 | H122460748 | O157:H7 | AAVVRE010000037.1 | UK | | Stx2 | Eru1 |
| EGE7625108.1 | H122960360 | O157:H7 | AAVVRI010000092.1 | UK | | Stx1 | lambdoid |
| EGE7644261.1 | H123780462 | O157:H7 | AAVVRM010000028.1 | UK | | Stx2 | Eru7 |
| EGE7696677.1 | H132360377 | O157:H7 | AAVVSB010000001.1 | UK | | Stx1 | lambdoid |
| EGE7750849.1 | H112340317 | O157:H7 | AAVVSL010000029.1 | UK | | Stx2 | Eru1 |
| EGE7793292.1 | H123200240 | O157:H7 | AAVVSS010000051.1 | UK | | Stx1 | lambdoid |
| EGE7868356.1 | H052720465 | O157:H7 | AAVVSU010000026.1 | UK | | Stx2 | Eru7 |
| EGE7880002.1 | H124600634 | O157:H7 | AAVVTC010000107.1 | UK | | Stx2 | Eru1 |
| EGE7946210.1 | H123980865 | O157:H7 | AAVVTR010000053.1 | UK | | Stx1 | lambdoid |
| EGE7955773.1 | H120540363 | O157:H7 | AAVVTU010000036.1 | UK | | Stx2 | Eru1 |

| | | | | | | | |
|---|---|---|---|---|---|---|---|
| EGE7978731.1 | H123760762 | O157:H7 | AAVVTZ010000010.1 | UK | | Stx2 | Eru7 |
| EGE8029056.1 | H121620525 | O157:H7 | AAVVTX010000050.1 | UK | | Stx2 | Eru5 |
| EGE8029099.1 | H121100178 | O157:H7 | AAVVTX010000053.1 | UK | | Stx1 | lambdoid |
| EGE8059842.1 | H104620471 | O157:H7 | AAVVUS010000061.1 | UK | | Stx1 | lambdoid |
| EGE8089196.1 | H063100370 | O157:H7 | AAVVUX010000016.1 | UK | | Stx2 | Eru7 |
| EGE8094777.1 | H122980845 | O157:H7 | AAVVUY010000106.1 | UK | | Stx1 | lambdoid |
| EGE8105221.1 | H113320446 | O157:H7 | AAVVVG010000064.1 | UK | | Stx1 | Eru1 |
| EGE8114167.1 | H135020660 | O157:H7 | AAVVVF010000015.1 | UK | | Stx2 | Eru7 |
| EGE8120612.1 | H043240557 | O157:H7 | AAVVVB010000025.1 | UK | | Stx2 | Eru7 |
| EGE8122814.1 | H121560360 | O157:H7 | AAVVVC010000003.1 | UK | | Stx2 | Eru1 |
| EGE8201356.1 | H123820316 | O157:H7 | AAVVVU010000025.1 | UK | | Stx2 | Eru7 |
| EGE8248496.1 | H121380674 | O157:H7 | AAVVWD010000068.1 | UK | | Stx1 | lambdoid |
| EGE8253777.1 | H133400671 | O157:H7 | AAVVWC010000072.1 | UK | | Stx2 | Eru7 |
| EGE8259112.1 | H113580158 | O157:H7 | AAVVWF010000051.1 | UK | | Stx1 | lambdoid |
| EGE8285006.1 | H122220222 | O157:H7 | AAVVWI010000053.1 | UK | | Stx1 | lambdoid |
| EGE8319594.1 | H132580296 | O157:H7 | AAVVWR010000124.1 | UK | | Stx2 | Eru2 |
| EGE8396302.1 | H123000125 | O157:H7 | AAVVXF010000060.1 | UK | | Stx1 | lambdoid |
| EGE8406126.1 | H103680174 | O157:H7 | AAVVXH010000123.1 | UK | | Stx2 | Eru7 |
| EGE8409932.1 | H123900440 | O157:H7 | AAVVXJ010000022.1 | UK | | Stx2 | Eru7 |
| EGE8411765.1 | H123120512 | O157:H7 | AAVVXJ010000073.1 | UK | | Stx1 | lambdoid |
| EGE8423942.1 | H125180252 | O157:H7 | AAVVXL010000004.1 | UK | | Stx1 | Eru4 |
| EGE8497699.1 | H094000312 | O157:H7 | AAVVXZ010000026.1 | UK | | Stx2 | Eru7 |
| EGE8529150.1 | H123880634 | O157:H7 | AAVVYD010000096.1 | UK | | Stx1 | lambdoid |
| EGE8534332.1 | H132480192 | O157:H7 | AAVVYG010000069.1 | UK | | Stx1 | lambdoid |
| EGE8565673.1 | H132080461 | O157:H7 | AAVVYL010000064.1 | UK | | Stx1 | lambdoid |
| EGE8879459.1 | H103460518 | O157:H7 | AAVVOT010000006.1 | UK | | Stx1 | lambdoid |
| EGE8936903.1 | H102980650 | O157:H7 | AAVVOS010000060.1 | UK | | Stx1 | lambdoid |
| KYR25636.1 | STEC 731 | O145:Hnt | LOFN01000117.1 | Netherlands | 2013 | Stx2 | Eru7 |
| KYR31563.1 | STEC 709 | O26:H11 | LOFM01000026.1 | Netherlands | 2013 | Stx2 | Eru7 |
| KYR32735.1 | STEC 709 | O26:H11 | LOFM01000001.1 | Netherlands | 2013 | Stx2 | Eru1 |
| KYR47546.1 | STEC 931 | O26:H11 | LOFS01000183.1 | Netherlands | 2013 | Stx2 | Eru1 |
| KYR50760.1 | STEC 886 | O91:H14 | LOFR01000041.1 | Netherlands | 2013 | Stx1 | Eru4 |
| KYR81241.1 | STEC 1188 | O91:H14 | LOFX01000035.1 | Netherlands | 2013 | Stx1 | Eru4 |
| KYS15770.1 | STEC 1293 | O26:H11 | LOGF01000197.1 | Netherlands | 2013 | Stx2 | Eru7 |
| KYS20709.1 | STEC 1299 | O128:H2 | LOGG01000109.1 | Netherlands | 2013 | Stx1 | Eru4 |
| KYS26052.1 | STEC1299 | O128:H2 | LOGG01000057.1 | Netherlands | 2013 | Stx2 | Eru6 |
| KYS37089.1 | STEC1375 | O185:H7 | LOGJ01000018.1 | Netherlands | 2013 | Stx2 | Eru7 |
| KYS42165.1 | STEC 1363 | O128:H2 | LOGI01000090.1 | Netherlands | 2013 | Stx2 | Eru6 |
| KYS44558.1 | STEC 1442 | O145:Hnt | LOGK01000083.1 | Netherlands | 2014 | Stx2 | Eru7 |
| KYS51717.1 | STEC 1465 | O128:H2 | LOGL01000097.1 | Netherlands | 2014 | Stx1 | Eru1 |
| KYS62574.1 | STEC 1500 | O76:H19 | LOGN01000031.1 | Netherlands | 2014 | Stx1 | Eru4 |

| | | | | | | | |
|---|---|---|---|---|---|---|---|
| KYS86001.1 | STEC1686 | O27:H30 | LOGT01000177.1 | Netherlands | 2014 | Stx2 | Eru10 |
| KYS86158.1 | STEC 1585 | O91:H14 | LOGR01000002.1 | Netherlands | 2014 | Stx1 | Eru4 |
| KYT16274.1 | STEC2193 | O103:H2 | LOGW01000035.1 | Netherlands | 2013 | Stx1 | lambdoid |
| KYT16329.1 | STEC 2236 | O113:H4 | LOGY01000073.1 | Netherlands | 2013 | Stx2 | lambdoid |
| KYT35599.1 | STEC 1506 | O5:H19 | LPWV01000121.1 | Netherlands | 2014 | Stx1 | Eru4 |
| KYT56146.1 | STEC 2363 | O91:H14 | LPWY01000046.1 | Netherlands | 2013 | Stx1 | Eru4 |
| KYT60009.1 | STEC 2419 | O103:H2 | LPWZ01000120.1 | Netherlands | 2013 | Stx1 | lambdoid |
| KYT71597.1 | STEC 2505 | O128:H2 | LPXB01000100.1 | Netherlands | 2013 | Stx2 | Eru6 |
| KYT71713.1 | STEC 2505 | O128:H2 | LPXB01000098.1 | Netherlands | 2013 | Stx1 | Eru4 |
| KYT73689.1 | STEC 2450 | O146:H21 | LPXA01000101.1 | Netherlands | 2013 | Stx1 | Eru4 |
| KYT80232.1 | STEC 2954 | O174:H8 | LPXE01000152.1 | Netherlands | 2013 | Stx1 | Eru4 |
| KYT81410.1 | STEC 2746 | O128ab:H2 | LPXD01000148.1 | Netherlands | 2013 | Stx2 | Eru6 |
| KYT86498.1 | STEC 2746 | O128ab:H2 | LPXD01000109.1 | Netherlands | 2013 | Stx1 | Eru4 |
| KYT97710.1 | STEC 2499 | O181:H49 | LOIE01000066.1 | Netherlands | 2013 | Stx1 | Eru7 |
| KYU10111.1 | STEC 2564 | O117:H7 | LOIG01000176.1 | Netherlands | 2013 | Stx1 | Eru1 |
| KYU17220.1 | STEC 2591 | O174:H2 | LOII01000032.1 | Netherlands | 2013 | Stx1 | Eru7 |
| KYU24994.1 | STEC 2633 | O146:H10 | LOIK01000066.1 | Netherlands | 2013 | Stx1 | Eru5 |
| KYU38540.1 | STEC 2764 | O174:H8 | LOIN01000127.1 | Netherlands | 2013 | Stx1 | Eru4 |
| KYU47608.1 | STEC 2788 | O91:H14 | LOIO01000033.1 | Netherlands | 2013 | Stx1 | Eru4 |
| KYU60947.1 | STEC 2841 | O nt:H20 | LOIQ01000003.1 | Netherlands | 2013 | Stx1 | Eru4 |
| KYU68016.1 | STEC 2894.1 | O1:H20 | LOIS01000032.1 | Netherlands | 2013 | Stx1 | Eru5 |
| KYU87735.1 | STEC2953 | O113:H4 | LOIW01000084.1 | Netherlands | 2016 | Stx2 | Eru10 |
| KYU88777.1 | STEC2953 | O113:H4 | LOIW01000078.1 | Netherlands | 2013 | Stx1 | Eru4 |
| KYU89986.1 | STEC 2980 | O128ab:H2 | LOIY01000194.1 | Netherlands | 2014 | Stx1 | Eru4 |
| KYU92507.1 | STEC 2962 | O91:H14 | LOIX01000034.1 | Netherlands | 2013 | Stx1 | Eru4 |
| KYV00368.1 | STEC2980 | O128ab:H2 | LOIY01000123.1 | Netherlands | 2014 | Stx2 | Eru6 |
| KYV06485.1 | STEC 3031 | O38:H26 | LOIZ01000129.1 | Netherlands | 2014 | Stx2 | Eru6 |
| KYV07180.1 | STEC 3031 | O38:H26 | LOIZ01000128.1 | Netherlands | 2014 | Stx1 | Eru4 |
| KYV16898.1 | STEC 2839 | O76:H19 | LOJB01000015.1 | Netherlands | 2013 | Stx1 | Eru4 |
| KYV23489.1 | STEC 2064 | O91:H14 | LOJC01000003.1 | Netherlands | 2013 | Stx1 | Eru4 |
| KYV25047.1 | STEC 2074 | O146:H21 | LOJD01000082.1 | Netherlands | 2013 | Stx1 | Eru4 |
| KYV25655.1 | STEC 66 | O165:H25 | LNFT01000172.1 | Netherlands | 2013 | Stx2 | Eru1 |
| KYV36935.1 | STEC 29 | O91:H14 | LNFU01000056.1 | Netherlands | 2013 | Stx1 | Eru4 |
| KYV47051.1 | STEC 196 | O91:H14 | LNZK01000067.1 | Netherlands | 2013 | Stx1 | Eru4 |
| KYV51123.1 | STEC 168 | O91:H14 | LNFV01000026.1 | Netherlands | 2013 | Stx1 | Eru4 |
| KYV59879.1 | STEC 200 | O174:H21 | LNZL01000033.1 | Netherlands | 2013 | Stx2 | Eru7 |
| KYV68949.1 | STEC 329 | O91:H14 | LOCT01000103.1 | Netherlands | 2013 | Stx1 | Eru4 |
| KYW17099.1 | STEC 559 | O182:H2 | LODC01000145.1 | Netherlands | 2013 | Stx1 | lambdoid |
| KYW31865.1 | STEC 565 | O121:H19 | LODE01000106.1 | Netherlands | 2013 | Stx2 | Eru2 |
| KYW38149.1 | STEC 645 | O91:H14 | LODG01000045.1 | Netherlands | 2013 | Stx1 | Eru4 |
| KYW59579.1 | STEC 3084 | O91:H14 | LPUJ01000040.1 | Netherlands | | Stx1 | Eru4 |

| | | | | | | | |
|---|---|---|---|---|---|---|---|
| KYW63559.1 | STEC 3087 | O91:H14 | LPUK01000020.1 | Netherlands | 2014 | Stx1 | Eru4 |
| KYW70309.1 | STEC 3106 | O91:H14 | LPUN01000125.1 | Netherlands | | Stx1 | Eru4 |
| KYW76135.1 | STEC 3098 | O174:H21 | LPUM01000015.1 | Netherlands | 2014 | Stx2 | Eru6 |
| MBH5127850.1 | 2018-439 | | JAEAMU010000071.1 | Switzerland | 2018 | Stx2 | Eru7 |
| MBH5152383.1 | 2017-353-1 | | JAEAMW010000071.1 | Switzerland | 2017 | Stx2 | Eru7 |
| MBH5157946.1 | S19-710-1 | | JAEAMY010000078.1 | Switzerland | 2019 | Stx2 | Eru7 |
| MBH5162990.1 | 2017-299-1 | | JAEAMX010000071.1 | Switzerland | 2017 | Stx2 | Eru7 |
| MBH5178762.1 | S19-30-1 | | JAEANC010000071.1 | Switzerland | 2019 | Stx2 | Eru7 |
| MBH5194740.1 | S19-18-1 | | JAEANE010000081.1 | Switzerland | 2019 | Stx2 | Eru7 |
| MBH5199905.1 | S19-101-1 | | JAEANF010000070.1 | Switzerland | 2019 | Stx2 | Eru7 |
| MBH5205094.1 | S18-73 | | JAEANH010000067.1 | Switzerland | 2018 | Stx2 | Eru7 |
| MBH5215298.1 | S18-9-1 | | JAEANG010000064.1 | Switzerland | 2018 | Stx2 | Eru7 |
| MBH5225647.1 | P17-291 | | JAEANK010000069.1 | Switzerland | 2017 | Stx2 | Eru7 |
| MBH5239801.1 | S18-168 | | JAEANJ010000068.1 | Switzerland | 2018 | Stx2 | Eru7 |
| MBI1440405.1 | STEC_UC4132 | | JACZHX010000017.1 | Italy | 2019 | Stx1 | Eru7 |
| MBI1456586.1 | STEC_UC4130 | | JACZHZ010000017.1 | Italy | | Stx1 | Eru7 |
| MBI1465333.1 | STEC_UC4131 | | JACZHY010000023.1 | Italy | | Stx1 | Eru7 |
| MBI1490075.1 | STEC_UC4128 | | JACZIB010000017.1 | Italy | | Stx1 | Eru7 |
| MBL6171648.1 | LSC6-3 | | JAETZC010000029.1 | Switzerland | 2020 | Stx2 | Eru10 |
| MBL6193510.1 | LSC1-7 | | JAETZA010000009.1 | Switzerland | 2020 | Stx1 | Eru4 |
| MBL6204510.1 | ATC7-7 | | JAETYU010000018.1 | Switzerland | | Stx1 | Eru4 |
| MBL6236395.1 | LSC1-58 | | JAETYZ010000032.1 | Switzerland | 2020 | Stx1 | Eru4 |
| MBL6291528.1 | LSC-5-20 | | JAETYY010000002.1 | Switzerland | 2020 | Stx1 | Eru4 |
| MBL6338105.1 | ATC36-6 | | JAETYQ010000045.1 | Switzerland | 2020 | Stx1 | Eru4 |
| MBL6358120.1 | ATC-4-67 | | JAETYK010000017.1 | Switzerland | 2020 | Stx1 | Eru1 |
| MBL6374145.1 | ATC-15-17 | | JAETYI010000026.1 | Switzerland | 2020 | Stx1 | Eru1 |
| MBL6378896.1 | ATB47-1 | | JAETYG010000030.1 | Switzerland | 2020 | Stx2 | lambdoid |
| MBL6401972.1 | ATB-14-66 | | JAETYA010000007.1 | Switzerland | 2020 | Stx1 | Eru4 |
| MBL6419043.1 | ATB-10-31 | | JAETXY010000022.1 | Switzerland | 2020 | Stx1 | Eru4 |
| MBL9224177.1 | STEC 559 | O182:H2 | LODC01000145.1 | Netherlands | 2013 | Stx1 | lambdoid |
| NP_309212.2 | Sakai | O157:H7 | NC_002695 | Japan | 2000 | Stx2 | Eru3 |
| NP_311017.2 | Escherichia coli str. Sakai | O157:H7 | NC_002695 | Japan | 2000 | Stx1 | lambdoid |
| NYR48883.1 | E7N18P5C8G | ONT:H28 | JACBWE010000011.1 | Portugal | 2019 | Stx1 | Eru7 |
| NYR49962.1 | E7N18P5C8G | ONT:H28 | JACBWE010000017.1 | Portugal | 2019 | Stx2 | Eru8 |
| NYR61457.1 | E7N15P4C10 | O116:H21 | JACBWG010000002.1 | Portugal | 2019 | Stx2 | Eru7 |
| NYR80058.1 | E7N6P4C8C | O29:H12 | JACBWK010000024.1 | Portugal | 2019 | Stx1 | Eru6 |
| NYR83788.1 | E7N6P4C8A | O29:H12 | JACBWL010000012.1 | Portugal | 2019 | Stx1 | Eru6 |
| NYS00014.1 | E7N3P7C8A | O150:H2 | JACBWM010000026.1 | Portugal | 2019 | Stx2 | Eru1 |
| NYS00969.1 | E7N3P7C8A | O150:H2 | JACBWM010000056.1 | Portugal | 2019 | Stx1 | lambdoid |
| PAT83787.1 | 536-9 | O26:H11 | MRVS01000020.1 | Israel | 2016 | Stx2 | Eru7 |
| PAT90662.1 | 479BS2 | O26:H11 | MRVR01000011.1 | Israel | 2016 | Stx2 | Eru7 |

| | | | | | | | |
|---|---|---|---|---|---|---|---|
| PAT95399.1 | 476-14 | O26:H11 | MRVU01000049.1 | Israel | 2016 | Stx2 | Eru7 |
| PAT99334.1 | 510-5 | O26:H11 | MRVV01000040.1 | Israel | 2016 | Stx2 | Eru7 |
| PAU07863.1 | 625C-4 | O26:H11 | MRVW01000016.1 | Israel | 2016 | Stx2 | Eru7 |
| PAU13589.1 | 514-2 | O174:H21 | MRVZ01000122.1 | Israel | 2016 | Stx2 | Eru7 |
| PAU30050.1 | 573-4 | O171:H29 | MRWA01000015.1 | Israel | 2016 | Stx2 | Eru7 |
| PRT57725.1 | St. Olav164 | | PVRW01000040.1 | Norway | | Stx2 | Eru7 |
| QIW91713.1 | Lys8385Vzw | O103:H11 | MT225100 | Japan | 2020 | Stx1 | Eru6 |
| QIW91773.1 | Lys19259Vzw | O157:H7 | MT225101 | Japan | 2020 | Stx2 | lambdoid |
| QKA14637.1 | NE 1092-2 | O157:H7 | NZ_CP038328.1 | USA | 2000 | Stx1 | Eru1 |
| QKA15101.1 | NE 1092-2 | O157:H7 | CP038328.1 | USA | 2000 | Stx2 | Eru2 |
| YP_001648917.1 | Enterobacteria phage Min27 | O157:H7 | NC_010237.1 | China | 2007 | Stx2 | Eru5 |
| YP_002274230.1 | Stx2-converting phage 1717, complete prophage genome | O157:H7 | NC_011357.1 | Canada | 2008 | Stx2 | Eru2 |
| YP_009226843.1 | Shigella phage 75/02 Stx | Shigella | NC_029120.1 | Hungary | 2013 | Stx1 | Eru4 |
| YP_009907967.1 | Escherichia phage SH2026Stx1 | O157:H7 | NC_049919.1 | USA | 2018 | Stx1 | Eru2 |
| YP_007001447.1 | TL-2011c | O103:H4 | NC_019442 | Norway | 2006 | Stx2 | Eru1 |
| AAD25430.1 | 933W | O157:H7 | NC_000924 | USA | 1982 | Stx2 | lambdoid |
| NP_040628.1 | lambda | | NC_001416 | | | | lambdoid |
| JAGEXB010000044 | BfR-EC-17679 | O36:H14 | JAGEXB010000044.1 | Switzerland | 2018 | Stx2 | Eru9 |
| KYU24344.1 | STEC 2595 | | LOIJ01000033.1 | Netherlands | 2013 | Stx2 | Eru11 |
| CCVP01000073 | FHI32 | | CCVP01000073.1 | Norway | 2009 | Stx1 | Eru12 |
| YP_794110.1 | Stx2-converting phage 86 | O86:H- | NC_008464.1 | Japan | | Stx2 | Eru3 |

**Supplemental Table S4**

Percent Identity Matrix (Clustal2.1) between clades of CI repressor sequences.

| | I | II | III | IV | V | VIa | VIb | VIIa | VIIIa |
|---|---|---|---|---|---|---|---|---|---|
| I | 100 | **51** | <25 | <25 | <25 | <25 | <25 | <25 | <25 |
| II | <25 | 100 | <25 | <25 | <25 | <25 | <25 | <25 | <25 |
| III | | | 100 | **60** | **32** | <25 | <25 | <25 | <25 |
| IV | | | | 100 | **42** | <25 | **35** | <25 | <25 |
| V | | | | | 100 | <25 | <25 | <25 | <25 |
| VIa | | | | | | 100 | **37** | <25 | <25 |
| VIb | | | | | | | 100 | <25 | <25 |
| VIIa | | | | | | | | 100 | <25 |
| VIIIa | | | | | | | | | 100 |
